# Isocitrate dehydrogenase 3b is required for spermiogenesis but dispensable for retinal degeneration

**DOI:** 10.1101/2022.02.09.479735

**Authors:** Siyan Zhu, Jiancheng Huang, Rong Xu, Yekai Wang, Yiming Wan, Rachel McNeel, Edward Parker, Douglas Kolson, Michelle Yam, Bradley Webb, Chen Zhao, Jianhai Du

## Abstract

Isocitrate dehydrogenase 3 (IDH3), a key enzyme in mitochondrial tricarboxylic acid (TCA) cycle, catalyzes the decarboxylation of isocitrate into α-ketoglutarate (αKG), converting NAD^+^ into NADH. We have found that IDH3 β subunit (IDH3B) is essential for IDH3 activity in multiple tissues. Loss of *Idh3b* in mice causes substantial accumulation of the isocitrate and its precursors in the TCA cycle, particularly in the testes, whereas the levels of the downstream metabolites remain unchanged or slightly increased. The *Idh3b*-knockout (KO) mice have normal visual function without retinal degeneration. However, the male KO mice are infertile. Loss of *Idh3b* causes energetic deficit and disrupts the biogenesis of acrosome and flagellum, resulting in spermiogenesis arrestment in sperm cells. Together, we demonstrate IDH3B controls its substrate levels in the TCA cycle and it is required for sperm mitochondrial metabolism and spermiogenesis, highlighting the importance of the tissue-specific function of the ubiquitous TCA cycle.

## Introduction

The mitochondrial tricarboxylic acid (TCA) cycle is a central metabolic pathway for energy production, macromolecule synthesis, cellular redox balance and cell signaling^1^. This cycle consists of a series of biochemical reactions to oxidize nutrients to generate ATP and intermediates for the synthesis of proteins, lipids and nucleotides. Beyond these classical functions, mitochondrial intermediates such as citrate, α-ketoglutarate (αKG) and succinate, are also important signaling molecules for post-translational modifications and epigenetic control of gene expression^1, 2^. The malfunction of the TCA cycle is implicated in a wide spectrum of disorders including neurodegenerative diseases, blindness, tumorigenesis, inflammation, obesity and male infertility^1, 3, 4, 5^.

Isocitrate dehydrogenase (IDH) that converts isocitrate into αKG, is a crucial control point for the TCA cycle. In mammalians, IDH exists in three isoforms: IDH1, 2 and 3. IDH1 and IDH2, localized in the cytosol and mitochondria respectively, catalyze the reversible interconversion of isocitrate and αKG using NADP(H)^+^ as cofactor. Their forward reaction (oxidative decarboxylation) and reverse reaction (reductive carboxylation) play critical roles in lipogenesis, redox homeostasis and cell proliferation^6, 7, 8, 9^. IDH3 is the classic TCA cycle enzyme, using NAD^+^ as its cofactor to catalyze the irreversible conversion of isocitrate into αKG and the reduction of NAD^+^ into NADH for mitochondrial respiration. IDH3 finely regulates the rate of the TCA cycle through substrate availability and allosteric regulation. IDH3 activity is stimulated by citrate, isocitrate, NAD^2+^, ADP, Mg^2+^ Mn^2+^ and Ca^2+^, but inhibited by NADH, αKG, ATP, and NADPH^6, 10, 11, 12^.

IDH3 is a heterotetramer composed of the αβ and αγ heterodimers. The α subunit is critical for the catalytic activity, but it requires the β and γ subunits for structural assembly and allosteric regulation. Without β or γ subunit, the α subunit alone has almost no activity^13, 14^. IDH3 subunits are encoded by *IDH3A, IDH3B,* and *IDH3G,* which express abundantly in mitochondria-rich tissues such as the heart, skeletal muscle and brain. Although ubiquitously expressed, human mutations or altered expression of IDH subunits are biased towards specific tissues in diseases. Mutations of *IDH3A* and *IDH3B* are identified in patients with inherited retinal degeneration, pseudocoloboma and epileptic encephalopathy^4, 15, 16, 17, 18^. *IDH3A* is associated with tumorigenesis and its expression is upregulated in patient samples from hepatocellular carcinoma and glioblastoma^19, 20^. Significantly reduced IDH3B is found in spermatozoa from patients with male infertility and poor sperm motility^5, 21, 22^. Loss of *Idh3a* in mice is embryonic lethal, but mice with *Idh3a* E229K mutation exhibits retinal degeneration^23^. In Drosophila melanogaster, *Idh3b*-mutant larval salivary gland cells fail to undergo mitochondrial fragmentation and oxidative phosphorylation, resulting in the death of these cells^24^. Although the enzyme structure, regulation and mutations have been well studied, how mutations or altered expression of IDH3 subunits impact mitochondrial metabolism to cause tissue-specific impairment *in vivo* remains elusive.

In this study, we investigate how the global loss of *Idh3b* in mice influences mitochondrial metabolism, visual function, retinal viability, and male fertility. We demonstrate that IDH3B is essential for IDH3 activity and TCA cycle intermediate metabolism. Surprisingly, testis mitochondria specifically rely on IDH3 for their mitochondrial metabolism and the loss of *Idh3b* leads to severe arrestment of spermiogenesis. However, IDH3B is dispensable for visual function and retinal viability. Our study provides fundamental information on the tissue-specific function and metabolic regulation of IDH3 and sheds new insight on understanding the role of the TCA cycle in retinal health, sperm development and male infertility.

## Results

### Global deletion of IDH3B reduces or abolishes IDH3 activity in the heart, brain, retina and testes

To determine the functions of IDH3B *in vivo,* we obtained *Idh3b* global (whole-body) KO mice from the Jackson Laboratory. The *Idh3b*-KO mice were generated by the injection of four gRNAs using CRISPR/Cas9 technology, resulting in 343 bp deletion in exons 2-4 (**Fig 1a**). Before validating the deletion of IDH3B, we evaluated the protein levels of IDH3 subunits in different tissues from the wild type (WT) mice by Western blot. Among the eight tissues we examined, the heart and brain were the two tissues with the most abundant IDH3B expression. To confirm the global deletion of *Idh3b* in mice, we chose four tissues for analyses: tissues with the most abundant IDH3B protein expression (heart and brain) and tissues with potential functional defects based on literature (retina and testis). Immunoblot results showed that IDH3B proteins level is similar between the heterozygous (HET) and the WT mice in all the four tissues, but undetectable in the homozygous (KO) mice, confirming that IDH3B is globally deleted in the KO mice (**Fig 1c**). To examine how IDH3 activity is affected when IDH3B is absent, we performed enzyme activity assays for retina, brain, heart and testis from WT, HET and KO mice. IDH3 activities in the retina, brain and testis from the KO mice were almost completely abolished; however, ~30% of IDH3 activity was preserved in the heart (**Fig 1d**). Since the HET mice had similar IDH3B protein level and IDH3 activities to the WT, we adapted the HET mice as the control group for most analyses to save animals and facilitate breeding. Taken together, we validated the global deletion of IDH3B in *Idh3b*-KO mice and showed that IDH3B is essential for IDH3 activity, especially in the retina, brain and testis.

**Figure 1.**
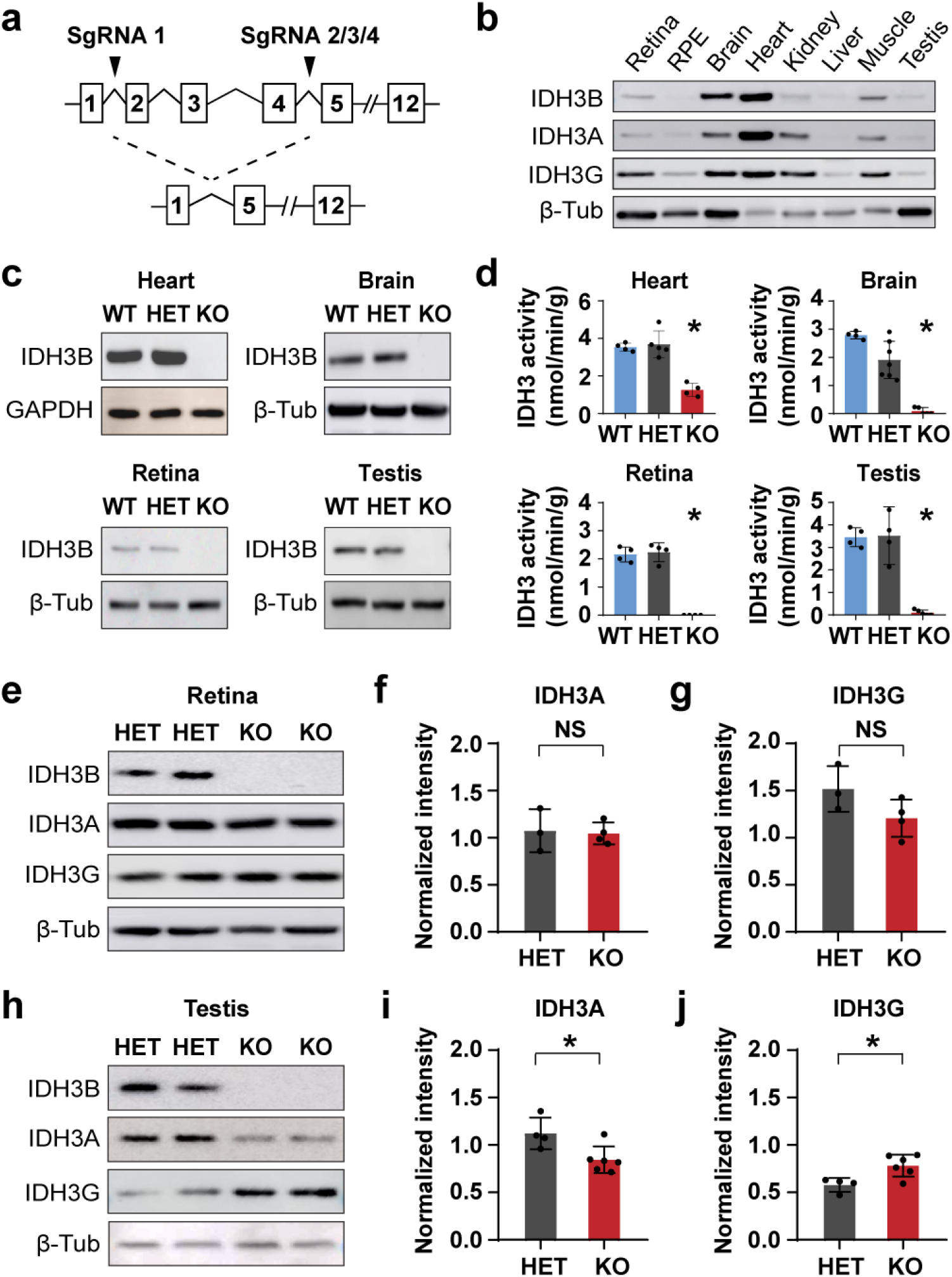
Generation of *Idh3b* knockout mice. **a** Schematic of *Idh3b*-KO mice with a deletion of exon 2-4 by CRISPR-Cas9. **b** Protein expression of IDH3A, IDH3B and IDH3G in retina, retinal pigmented epithelium (RPE), Brain, Heart, Kidney, Liver, Muscle, and Testis from WT mice at P80. **c** Western blot analysis for IDH3B in heart, brain, retina and testis from WT, HET and KO mice. GAPDH was used as loading control for heart and β-Tubulin (β-Tub) was used as loading control for brain, retina and testis. Repeated for three times. **d** Enzymatic activity analysis of IDH3 in heart, brain, retina and testis from WT, HET and KO mice at P50. N ≥ 4. *P < 0.05 KO versus WT or HET. **e-g** Western blot analysis of IDH3A and IDH3G expression in retinas from mice at P200. **e.** Representative Western blot image. **f-g** Semi-quantification of blots by normalizing with the loading control β-Tubulin. N ≥ 3 *P < 0.05. **h-j** Western blot analysis of IDH3A and IDH3G expression in testis from mice at P50. e. Representative Western blot image. f-g Semi-quantification of blots. N ≥ 4 *P < 0.05. (Data are all represented as mean ± SD. test.)

### Loss of Idh3b changes the expression levels of IDH3A and IDH3G subunits in testis but not retina

To investigate whether the loss of *Idh3b* altered the expression of other IDH3 subunits, we quantified the protein expression of IDH3A and IDH3G in the retina and testis. In the retina, densitometry data showed no significant difference in IDH3A and IDH3B protein levels between HET and KO (**Fig 1e-g)**. However, in the testis, IDH3A was reduced by 75% (P< 0.05), while IDH3G was increased by 35% (P< 0.05) in the KO mice (**Fig 1h-j)**. Overall, our results suggest that IDH3B is required for the proper expression of other IDH3 subunits in the testis but not the retina.

### Idh3b knockout mice have normal growth and vision without retinal degeneration

The *Idh3b*-KO mice showed normal appearance, hair color, and body weight (**Fig S1a-b**). There was no difference in the tissues/body weight in major organs including heart, kidney and liver between the KO and HET mice, suggesting that *Idh3b*-KO mice have normal body growth (**Fig S1c-e**). To investigate whether the KO mice have retinal degeneration, we examined the visual function by electroretinogram (ERG), and retinal morphology by optical coherence tomography (OCT) *in vivo* and Hematoxylin and Eosin (H&E) staining of eye sections *in vitro.* ERG responses from postnatal day 115 (P115) to P250 progressively declined with aging at the same rate between the KO and HET mice (**Fig S2a-d).** At P250, there was no significant changes in scotopic and photopic a-wave and b-wave at all light intensities in KO mice, compared with the HET (**Fig 2a-d**). OCT showed no difference in the thickness of each retinal layer between the KO and HET mice at the age of P200 (**Fig S2e-f**). Quantification of the thickness of the photoreceptor layer (outer segment and inner segment) and the number of nuclei in outer nuclei layer from H&E stained eye sections confirmed that *Idh3b*-KO mice had normal retinal morphology at P250 (**Fig 2e-h**). Together, these results demonstrate that *Idh3b*-KO mice have normal vision and retinal morphology without retinal degeneration.

**Figure 2.**
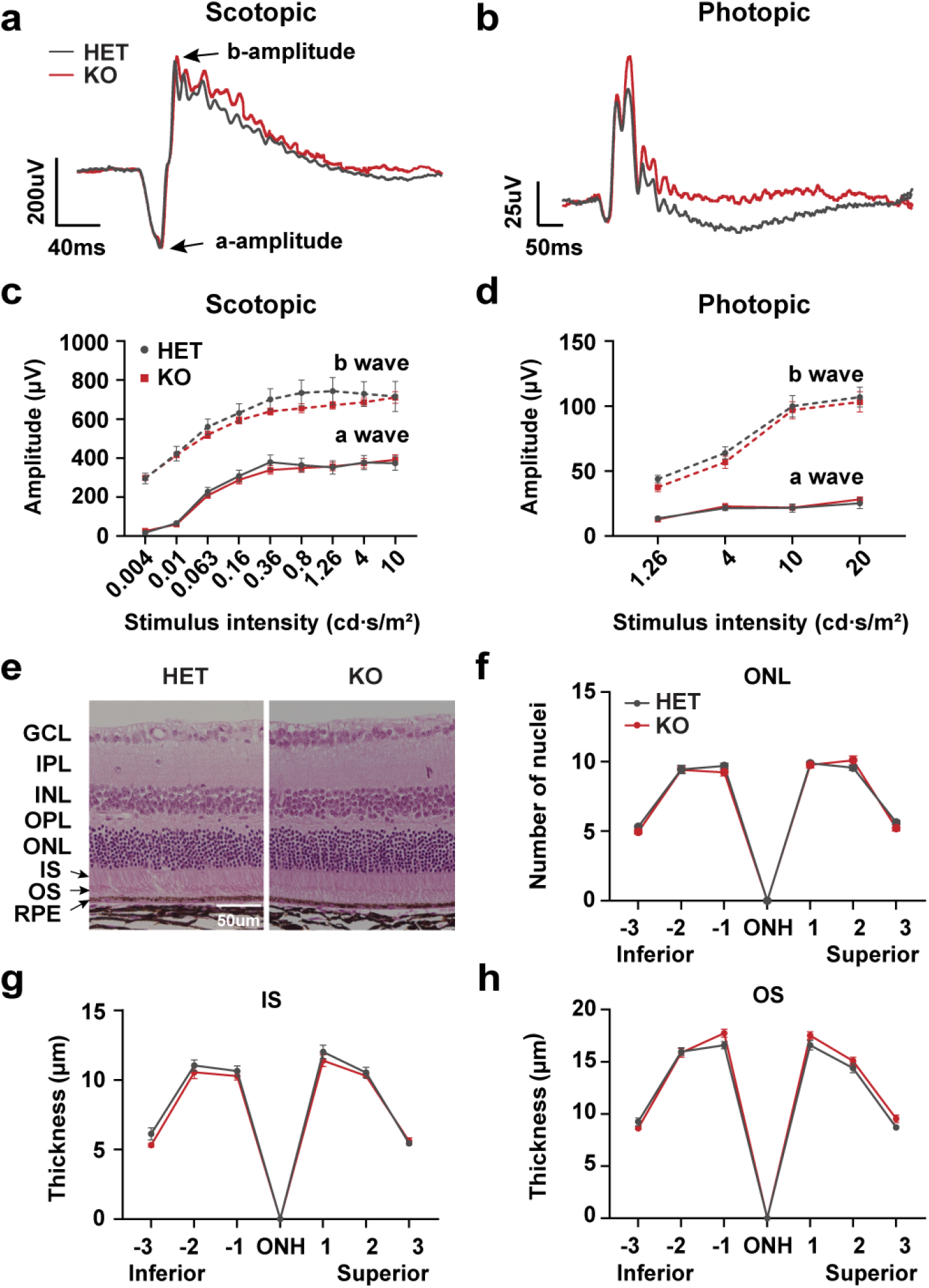
Loss of *Idh3b* did not affect visual function and retinal health in mice. **a-b** Representative scotopic (0.8 cd*s/m2) and photopic (10 cd*s/m2) ERGs of *Idh3b* KO and *Idh3b* HET (control) at P250. N ≥ 4. **c-d** Scotopic ERG and Photopic ERG analysis under different light intensities for HET and KO mice at P250. N ≥ 4. **e** Representative retinal H&E images from HET and KO mice at P250. **f** Number of nuclei in outer nuclei layer (ONL) at different positions of the eye. N ≥ 3 **g-h** Thickness of inner segment (IS) and outer segment (OS) at different positions of the eye. N ≥ 3. (Data are all represented as mean ± SD. t test.)

### The substrates of the IDH3 reaction are accumulated, but the products remain unchanged in Idh3b KO retinas

IDH3 irreversibly converts isocitrate from citrate and aconitate into αKG in the TCA cycle (**Fig 3a**). To examine how *Idh3b* ablation affects mitochondrial intermediary metabolism, we measured the abundance of TCA cycle metabolites from freshly isolated mouse retinas using Gas Chromatography Mass Spectrometry (GC MS). As expected, metabolites from upstream reactions of IDH3, including citrate, isocitrate and aconitate, were accumulated about 2-fold in retinas from the KO mice (**Fig 3b**). However, αKG and its downstream metabolites including glutamate, succinate, fumarate, malate and aspartate, remained unchanged (**Fig 3b**). These TCA cycle metabolites can be generated from different nutrient sources such as glucose, fatty acids and amino acids. To trace the changes of metabolites from the same nutrient source, we incubated freshly isolated retinas with ^13^C glucose and analyzed ^13^C glucose-labeled metabolites by GC MS as previously reported^25, 26^. Our results showed the abundance of ^13^C glucose-labeled glycolytic intermediates including phosphoenolpyruvate, pyruvate and lactate were similar between the HET and KO retinas. For TCA cycle intermediates, ^13^C glucose labeled citrate, isocitrate and aconitate increased 2~3 fold in the KO retinas, while other intermediates remained unchanged (**Fig 3c**). These results suggest IDH3B is critical for the levels of its substrates but not products. To examine whether another two IDH isoenzymes (IDH1 and IDH2) compensated for the loss of IDH3B, we measured their protein expression and enzyme activity from the isolated retinas. There was no significant difference in either IDH1 or IDH2 expression level between HET and KO (**Fig 3d-f**). The retinas from HET and KO showed similar NADP-dependent IDH activity from the IDH1 and IDH2 (**Fig 3g**). Overall, these results indicate IDH3B is essential for mitochondrial isocitrate oxidation in the retina and for maintaining the levels of TCA cycle metabolites.

**Figure 3.**
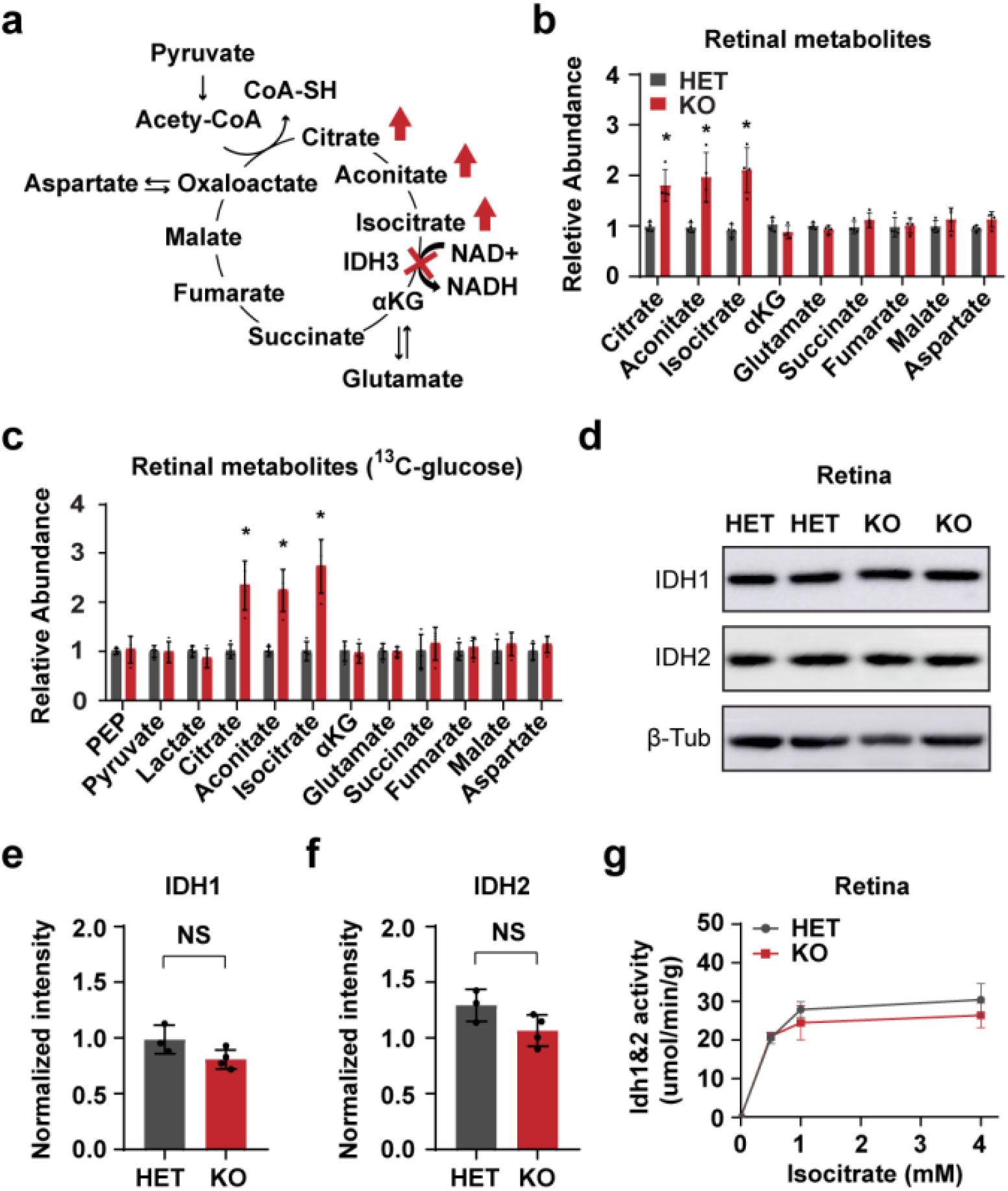
Loss of *Idh3b* disrupted cellular metabolism in the retina. **a** Schematic of TCA cycle metabolism. **b** Fold change of TCA cycle metabolites in retina from KO mice. N = 4 *P < 0.05. **c** Fold change of ^13^C-glucose-derived metabolites in the retina from KO mice. N ≥ 4 *P < 0.05. **d-f** Western blot analysis of IDH1 and IDH2 in the retina from mice at P200. d Representative Western blot image. **e-f** Semi-quantification of IDH1 and IDH2 level in retina. β-Tubulin (β-Tub) was used as loading control. N ≥ 3. **g** Enzymatic activity of IDH1&2 under different isocitrate concentrations in the retina from HET and KO mice. N ≥ 4.

### Idh3b KO male mice are sterile with no matured sperms in the epididymis

During the generation of *Idh3b*-KO mice, we observed that male KO mice could not generate pups when bred with either HET females or WT females for up to six months. However, the breeding of KO females with HET males was normal. We performed a gross examination of isolated testes, and found the size and weight of testes is similar between the HET and KO (**Fig S3a-b**). To ask whether the sperms were normal in *Idh3b*-KO mice, we dissected the cauda epididymis (where matured sperm are stored), live-imaged and counted the number of sperm cells under the light microscope. All sperm cells from the HET mice were motile with normal head-tail morphology **(Fig 4a, Supplemental Video 1)**. However, the KO mice had nearly no mature spermatozoa and showed a large accumulation of immotile round sperm cells. **(Fig 4b, Supplemental Video 2)**. By counting the sperm cells, we found both WT and HET mice had ~ 6 million healthy and mature spermatozoa, whereas the KO mice had no mature spermatozoa except ~ 0.8 million round sperm cells **(Fig 4c-d)**. Histological analysis of the cauda epididymis using H&E staining confirmed that epididymal ducts in the KO mice were filled with immature round sperm cells **(Fig 4e-f).** These immature cells exhibited multiple defects including enlarged and vacuolated cytoplasm, vacuolated nucleus, multinucleation and fragmentation (**Fig 4e’-f’**). Overall, we found that *Idh3b*-KO male mice were infertile without mature spermatozoa, indicating that sperm development is impaired in the *Idh3b*-KO mice.

**Figure 4.**
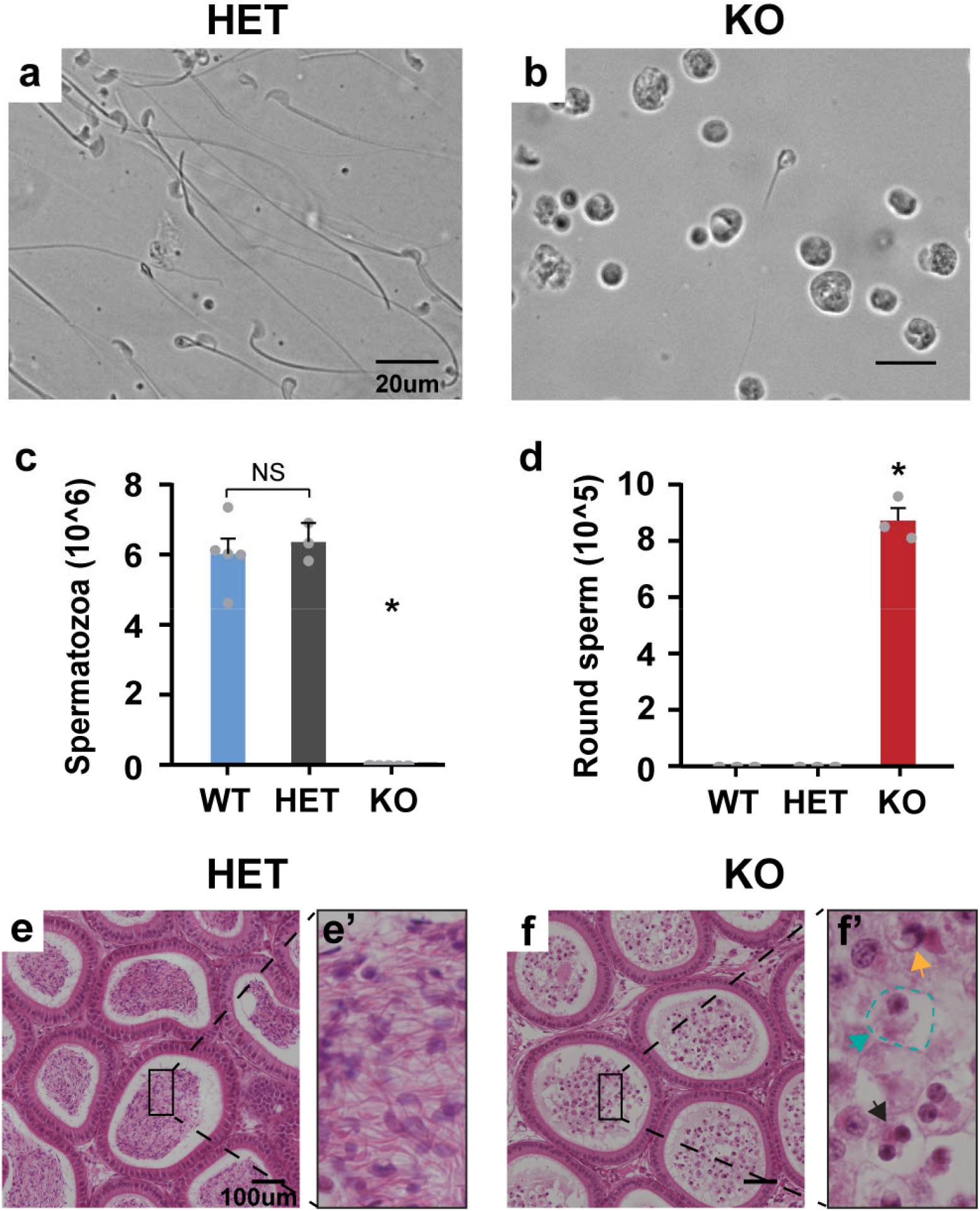
*Idh3b* male KO mice are sterile and lack of the ability to produce mature spermatozoa. **a-b** Representative images of sperm cells obtained from the cauda epididymis of HET and KO mice at P50. **c-d** Quantifications of mature spermatozoa and round sperm cells in the cauda epididymis. N ≥ 3 *P < 0.05 versus WT or HET. Data are represented as mean ± SD. t test. **e-f** H&E analysis of cauda epididymis in HET and KO mice at P50. **e’-f’** Magnified H&E images (green arrow: enlarged and vacuolated cytoplasm, yellow arrow: vacuolated nucleus, black arrow: fragmented nucleus).

### Spermiogenesis is arrested in Idh3b KO mice

Sperm development occurs in seminiferous tubules of the testis and consists of three stages: mitosis, meiosis and spermiogenesis. To investigate how *Idh3b* ablation influenced sperm development, we stained cross-sectioned testis and quantified different cell populations in seminiferous tubules. There was no difference in the number of cells in the early developmental stages, including spermatogonia, pachytene, leptotene/zygotene and spermatid, between the HET and KO mice **(Fig 5a-c)**, suggesting that sperm cells have normal development in mitosis and meiosis in the KO mice. However, the number of mature spermatozoa was reduced ~5 fold in the KO testis, indicating a spermiogenesis arrestment in *Idh3b*-KO mice **(Fig 5c)**. Mouse spermiogenesis can be sub-divided into 16 steps based on the morphological changes of the acrosome, a cap-like organelle attached to the nucleus (**Fig S4a**). To define the impaired steps in spermiogenesis in the KO mice, we stained the glycoprotein in acrosome by Periodic Acid Schiff (PAS). We found spermatids at the same differential step orderly encircled the seminiferous tubules in HET testis but were disorganized in the KO testis (**Figure S4b**). Detached or malformed acrosomes were observed in KO but not HET in steps 2-7 (**Figure 5d-i**). HET mice developed a normal oval shape nucleus starting from step 8 (**Fig 5h, j, l**). However, the shapes of the nucleus in the KO testis were irregular with round heads. Additionally, flagella (the tails), another important feature in mature spermatozoa, were abundant in the HET but scarce in KO mice (**Fig S4b**). These results indicate that loss of *Idh3b* results in spermiogenesis arrestment.

**Figure 5.**
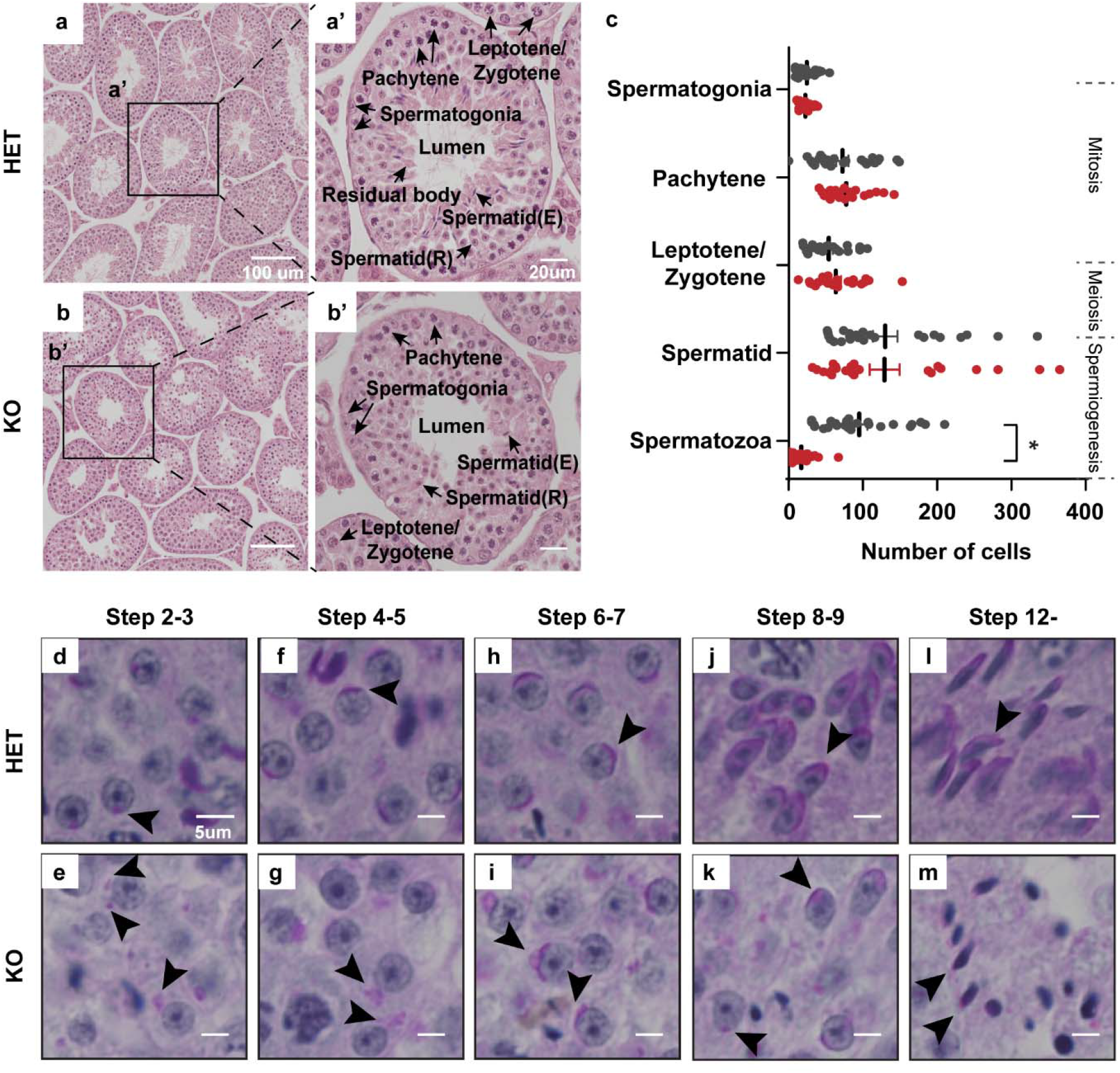
IDH3B is essential for spermiogenesis. **a-b** Representative H&E images for testicular sections from mice at P50. a’-b’ Late-stage testicular tubule with mature spermatozoa. spermatid(R): round spermatid, spermatid(E): Elongated spermatid. **c** Quantification of different sperm cell populations in testis. N of tubules = 20 *P <0.05. Data are represented as mean ± SD. t test. **d-m** Periodic acid-Schiff stain (PAS) staining for staging sperm differentiation in testes. Arrow heads pointed to the acrosome.

### Loss of IDH3B disrupts acrosome biogenesis

The acrosome is formed by organelles-derived (particularly Golgi) proacrosomal vesicles that traffick, fuse and anchor to the nuclear surface during spermiogenesis. Acrosome biogenesis can be divided into four phases: Golgi (steps 1-3), Cap (steps 4-7), Acrosome (steps 8-12) and Maturation (steps 13-16) phases (**Fig S4a**). To visualize acrosome biogenesis, we stained the acrosome, nucleus and Golgi with PNA, DAPI and GM130. In steps 2-3 (Golgi phase), a single and sphere acrosomal granule is attached to the nucleus in the HET testis, but detached in the KO with or without misoriented Golgi, indicating a compromised vesicle trafficking and anchoring in the KO mice **(Fig 6a-b)**. In steps 4-5 (Cap phase), proacrosomal granules displayed abnormal shapes and failed to transform into a cap shape in the KO testis **(Fig 6c-f)**. In steps 6-7 (Cap phase), cap-shaped proacrosomal vesicles were orderly oriented towards the nucleus in the HET but the proacrosomal vesicles were scattered to float around the nucleus in the KO testis, indicating vesicle trafficking and fusion impairments. In the later steps (Acrosome and Maturation Phases), the HET mice developed elongated mature acrosomes, whereas the KO acrosomes were fragmented and disoriented. **(Fig 6g-j)**. To confirm these findings, we examined the ultrastructure of acrosome biogenesis using the transmission electron microscope (TEM). Consistently, we found multiple defects in acrosome biogenesis in the KO. These defects included: 1) Golgi misorientation and acrosome detachment in Golgi phase, 2) no acrosome or eccentric and dilated acrosome granules in Cap phase, 3) no acrosome or immature acrosome with mislocalized acrosomal granule in Acrosome phase, and 4) no acrosome or detached acrosome in Maturation phase **(Fig 6l-v)**. To further evaluate these defects, we measured the expression of key proteins in acrosome biogenesis, including Heat Shock Protein 90 Beta Family Member 1 (HSP90B1), Golgi Associated PDZ and Coiled-Coil Motif Containing (GOPC), and Acrosomal Vesicle Protein 1 (ACRV1) **(Fig 6w-x)**. HSP90B1, an ER protein required for vesicle formation ^27^, was similar between the KO and HET. GOPC, which is required for vesicle fusion, was also maintained with the same amount in the KO ^28^. Remarkably, ACRV1, an acrosomal matrix protein arising during acrosome biogenesis, was significantly decreased in testis from the KO mice^29^. Together, these results strongly support that IDH3B is essential for acrosome biogenesis.

**Figure 6.**
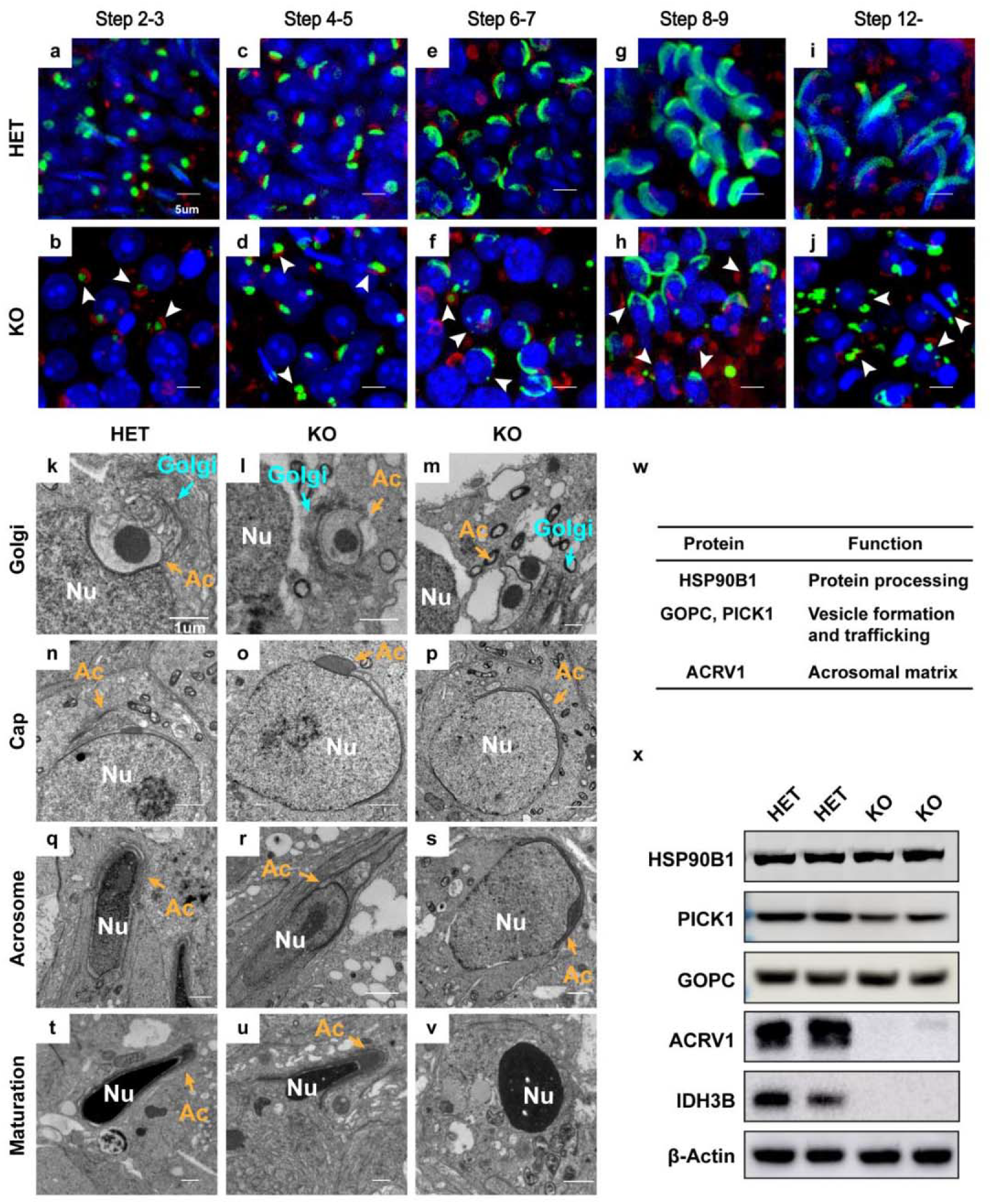
Acrosome biogenesis defects in *Idh3b* KO mice. **a-j** Representative images for immunofluorescent staining of acrosome and Golgi at different stages in testicular tubular sections from mice at P50. Red: GM130, Blue: DAPI, Green: PNA. Arrowhead: defected acrosome. **k-v** Ultrastructure analysis of acrosome biogenesis at Golgi, Cap, Acrosome and Maturation stages. Blue arrows: Golgi, yellow arrows: Acrosome (Ac), Nucleus (Nu). **w** Table of proteins essential for acrosome biogenesis. Hsp90b1: heat shock protein 90 beta family member 1, GOPC: Golgi associated PDZ and coiled-coil motif containing, PICK1: Protein interacting with PRKCA 1, ACRV1: Acrosomal vesicle protein 1. **x** Western blot analysis of listed proteins in testis from HET and KO mice at P50.

### Idh3b is required for flagellum assembly

Flagellum biogenesis is another hallmark for spermiogenesis. In KO testis, we found much fewer residual bodies and flagella in the lumen of testicular tubules **(Fig 5b’)**. To investigate how flagellum formation is affected, we stained axoneme, the core structure of flagellum, with the antibody against acetyl-α-Tubulin. In testicular tubules containing late-step spermatids, we found that the distal-ends of flagella were well oriented toward the lumen in the HET but were misoriented and disorganized in the KO mice **(Fig 7a-b)**. The mitochondrial sheath is an important structural component, where mitochondria align with axoneme in the flagellum. Fluorescent staining of mitochondria using Mito-tracker showed substantially reduced signals in the KO sections, suggesting that the mitochondrial sheath is disrupted. **(Fig 7e-f)**. Fibrous sheath and outer dense fibers are crucial for the assembly and integrity of the flagellum. To quantitatively study the structural changes in the flagellum, we performed Western blot with antibodies against Acetyl-tubulin (axoneme marker), A-Kinase Anchoring Protein 4 (AKAP82) (fibrous sheath marker), and Outer Dense Fiber 2 (ODF2) (outer dense fiber marker). Our results showed that all these proteins were significantly reduced, especially AKAP82, which was almost undetectable in the KO testis **(Fig 7g-h)**. To validate these findings, we examined the ultrastructure of flagellum components using TEM. We found structural defects in all three pieces in the flagellum from the KO testis. The mitochondrial sheath disappeared and multi-flagella were formed in the mid piece **(Fig 7i-k)**. In the principal piece, there were abnormal axonemes and reduced longitudinal columns **(Fig 7l-n)**. In the end piece, the plasma membrane was irregular and disorganized, which is a notable atypical feature of defected flagellum **(Fig 7o-q)**. Together, our results suggest that IDH3B is critical for maintaining intact flagellar structure including axoneme, mitochondrial sheath, fibrous sheath and outer dense fiber.

**Figure 7.**
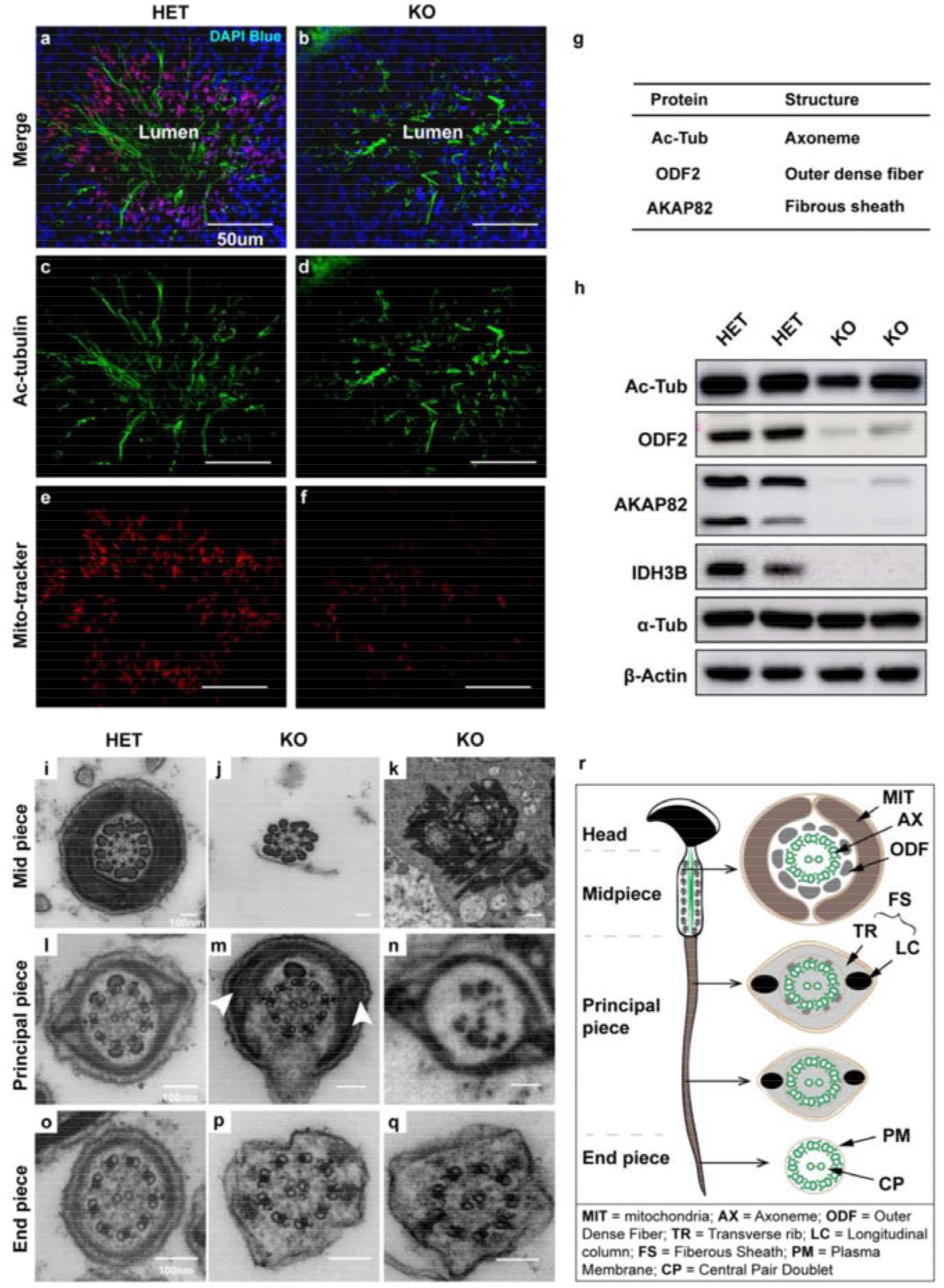
Flagellum biogenesis defects in *Idh3b* KO mice. **a-f** Immunofluorescent staining of axoneme and mitochondria in late-stage differentiating testicular tubule from mice at P50. Red: Mito-tracker, Blue: DAPI, Green: acetyl-α-tubulin. **g** Table of proteins as structural components for flagellum integrity. Actubulin: acetyl-α-tubulin, ODF2: Outer Dense Fiber of Sperm Tails 2, AKAP82: A-kinase Anchor Protein 82Kd. **h** Western blot analysis of listed proteins in testis from HET and KO mice at P50. **i-q** Ultrastructure analysis of different pieces of flagellum from mice at P50. Arrow heads pointed towards the longitudinal column. **r** A schematic of flagellum structures.

### Idh3b is specifically required for testis mitochondrial metabolism

To test how the loss of IDh3b influences testis metabolism, we firstly measured TCA cycle metabolites as done previously with the retinas using GC MS. Surprisingly, metabolites upstream of the IDH3 reaction, including citrate, isocitrate and aconitate, increased 10~20-fold in testes from the KO, compared to the Control **(Fig 8a)**. However, the metabolites downstream of the IDH3 reaction were either slightly increased or unchanged (**Fig 8a**). To ask whether this massive accumulation of precursors was specific to the testes, we analyzed TCA cycle intermediates from hearts where IDH3b protein is most abundant. Similar to the retina, metabolites upstream of the IDH3 reaction in the IDH3B KO hearts increased only 2~3 fold (**Fig 8b)**. NADP-dependent IDH1 and IDH2 can also oxidize isocitrate. There was no significant difference in protein expression and enzyme activities of IDH1/2 between the HET and KO testes (**Fig S5 a-d**). IDH1/2 activities increased with isocitrate and reached the maximal activities at ~2mM. These results suggest that the massive accumulation of substrates probably exceeds the maximal rates of IDH1/2 for full compensation. To examine the impact of *Idh3b*-KO on the metabolome, we performed targeted metabolomics covering 206 metabolites in major metabolic pathways in glucose, amino acid and nucleotide metabolism (**Table S3**). Multivariant analysis showed that the scores plot of metabolites from the KO were well separated from the HET, demonstrating the loss of *Idh3b* significantly alters testis metabolome **(Fig 8c)**. Citrate and isocitrate were not in the targeted metabolomics, but we found citraconic acid (formed by the heating of citrate) and aconitate were substantially increased in the KO samples, consistent with the GC MS results. Notably, multiple acyl-carnitines, acetylated amino acids, the ketone body 3-hydroxybutyrate, and phosphocreatine were increased in the KO testes, suggesting there is an energetic deficit **(Fig 8d-e)**. Consistent with these results, ATP, NADH and UDP were significantly decreased in testes from the KO mice **(Fig 8d-e)**. However, the heart metabolome between the HET and KO were mostly overlapping, and high energy metabolites such as ATP, NADH and phosphocreatine were normal in the KO hearts (**Fig 8f-g**). Most acyl-carnitines were normal or decreased rather than accumulated in the KO hearts. These results suggest that IDH3b is specifically required in testis for mitochondrial metabolism.

**Figure 8.**
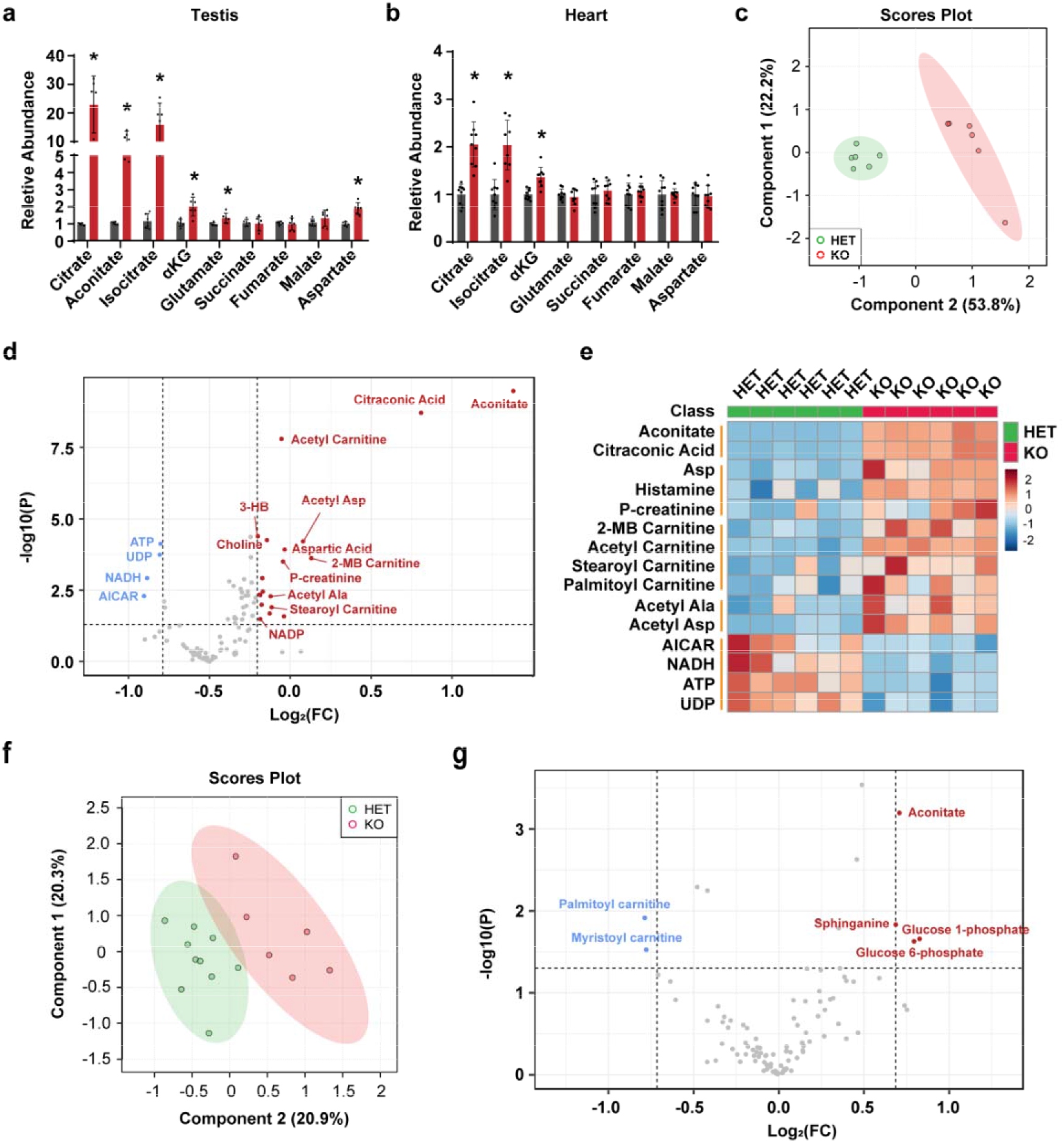
Loss of *Idh3b* dramatically disrupted cellular metabolism in testis but not the heart. **a** Fold change of TCA cycle metabolites in testis from KO mice at P50. N ≥ 6 *P < 0.05. **b** Fold change of TCA cycle metabolites in the heart from KO mice at P50. N ≥ 8 *P < 0.05. **c** Multivariant analysis of metabolomic study for testis by LC MS. N = 6. **d** Volcano plot of metabolites from the testes by LC MS. N ≥ 6, Fold change ≥ 1.5, P < 0.05 **e** Heatmap for top 15 changed metabolites. N=6, Fold change ≥ 1.5, P < 0.05 **f** Multivariant analysis of metabolites from the heart. N ≥ 8. **g** Volcano plot of metabolites from the heart by LC MS. N ≥ 8, Fold change ≥ 1.5, P < 0.05. (Data are all represented as mean ± SD. t test.)

## Discussion

In this study, we have found that IDH3B is essential for IDH3 activity in the retina, heart, brain, and testis. As a TCA cycle enzyme, IDH3 is important for maintaining the levels of citrate, isocitrate and aconitate, especially in the testis. IDH3B is dispensable for visual function and photoreceptor survival. However, it is critical for testis metabolism and sperm development. The testis specifically requires IDH3B to maintain its mitochondrial bioenergetics and acrosome and flagella biogenesis during spermiogenesis.

The full activity of IDH3 requires the assembly and cooperation of αβ and αγ heterodimers. *In vitro* biochemical reconstitution of subunits shows that, α, β or γ subunit alone is nearly inactive, but αγ heterodimer exhibits activity that weaker than the fully assembled enzyme^13^. IDH3B plays a critical structural role in assembling the holoenzyme in the forms of heterotetramer or heterooctamer^14, 30^. We found IDH3B ablation almost completely abolishes IDH3 activity in different tissues **(Fig 1d)**, indicating that the β subunit is important for the assembly or stability of αγ heterodimer *in vivo.* These results are consistent with the finding in the larval salivary gland cells in Drosophila melanogaster that the presumable *Idh3b* null allele blocks mitochondrial metabolism^24^. However, it is still unclear why the loss of *Idh3b* alters the protein expression of other subunits in the testis but not the retina. It is possible that IDH3 subunits in the testis have their specific interactions and/or post-translational regulation that govern their protein stability, which has been discovered for protein subunits in other systems^31, 32^

All the *Idh3-KO* tissues that we measured showed an accumulation of precursors (citrate, isocitrate and aconitate), particularly in the testis, but the level of the products (αKG and succinate) was unchanged or slightly increased. Under normal conditions, citrate from glucose, lipids or amino acids is rapidly utilized through 1) TCA cycle by aconitase and IDH3, 2) IDH1 and IDH2, and 3) ATP citrate lyase to be cleaved into acetyl-CoA as acetyl group donor and oxaloacetate as a substrate for aspartate and malate. In addition to IDHs, αKG can also be produced from glutamate through multiple transaminases and glutamate dehydrogenase^33^. It remains to be understood how these pathways compensate to maintain the levels of the products of the IDH3 reaction. The increase of aspartate, acyl-carnitines and acetylated amino acids in the KO testes further suggest the compensation of these pathways, but these compensations are incapable of maintaining the levels of NADH, ATP and other high energy metabolites. Based on our findings, we speculate that IDH3 may be the primary dehydrogenase for mitochondrial NADH production in the testis.

IDH3 mutations have been correlated with inherited retinal degeneration^4, 15, 16, 17^. Patients with *IDH3A* mutations showed an early onset (1-11 years old) of retinal degeneration and patients with *IDH3B* mutation showed a late onset (30-50 years old) of retinal degeneration^4, 15, 16, 17^. The loss of *Idh3a* in mice is embryonically lethal and the mouse *Idh3a* E229K mutation leads to an early and rapid retinal degeneration^23^. However, the loss of *Idh3b* in mice through an 8-bp deletion in exon 5 shows no retinal degeneration at P180^23^. This report is consistent with our findings that *Idh3b*-KO mice have normal vision up to P250, which is equivalent to 30 to 40 years age in humans. It is possible that human *IDH3B* mutants may have an unknown malfunction such as gain of function and non-metabolic function to cause retinal degeneration, which has been identified in mutations of other enzymes. For example, the mutations of inosine monophosphate dehydrogenase 1 *(IMPDH1),* a key enzyme in *de novo* nucleotide synthesis, cause severe retinal degeneration in patients, but one-year-old *Impdh1* knockout mice maintain normal retinal structure^34^. The *Impdh1* mutants are proposed to reduce binding to nucleotide and association with polyribosomes^32^. Another example is *IDH2.* Its mutations can gain a new function to generate 2-hydroxyglutarate, an oncometabolite in tumorigenesis^35^. Interestingly, IDH3A has been reported to localize in the nucleus and interact with transcriptional factors to promote tumor growth^19^. It is unknown whether IDH3B or its disease mutants moonlight similar other activities. Future study of the function of the mutant IDH3B proteins is warranted to elucidate the conundrum of retinal phenotypes in humans.

Why is IDH3B or IDH3 crucial for sperm development? There is a shift reliance of metabolism from glycolysis to TCA cycle during sperm development^36^ The aerobic oxidation of lactate, glucose, pyruvate and fatty acids through the TCA cycle is required to meet the high energetic demands in the maturation of spermatids ^37, 38, 39^. The TCA cycle enzymes are generally regarded as ubiquitous in nucleated cells for their housekeeping functions. However, the deficiencies of these enzymes including fumarase, succinate dehydrogenase, and αKG dehydrogenase show tissue-specific impairment^40, 41, 42^. Our findings demonstrate that IDH3B is specifically required for IDH3 activity, mitochondrial metabolism, and spermiogenesis. IDH3B proteins are under-expressed in low motile spermatozoa or spermatozoa from infertile men with varicocele^21^. IDH inhibitor can block sperm capacitation and inhibit acrosome reaction in boar spermatozoa^43^. Intriguingly, Gossypol, a natural antifertility agent for males, causes specific mitochondrial damage in the tails of spermatozoa^44, 45^, which has a similar phenotype to the *Idh3b*-KO mice. The block of IDH3 may be attributed to its induction of male infertility, as Gossypol has been a known inhibitor of NAD-linked dehydrogenase including IDH3^46^. The environmental contaminant Tributyltin, known to impair sperm maturation in the vertebrate^47, 48^, is also a sensitive inhibitor of IDH3^49^.

The IDH3 reaction controls the levels of key metabolites (citrate, NADH and ATP), which are essential for biosynthesis, cell signaling and bioenergetics during sperm development^50, 51, 52^. Besides the TCA cycle, citrate is an important source of acetyl-CoA, a substrate for *de novo* lipid synthesis and protein acetylation. During sperm differentiation, different compositions of lipid species are required to generate lipid-rich organelles like acrosome and flagellum^53^. Compared with immature spermatids, mature sperms have high amounts of polyunsaturated fatty acids (PUFA)^53^. The disruption of key enzymes in PUFA synthesis such as acyl-CoA synthetase 6 and delta-6 desaturase can arrest spermiogenesis, resulting in male infertility ^54, 55, 56, 57^ with similar phenotypes to the *Idh3b*-KO mice. Additionally, NADH is a required cofactor for key desaturases including delta-6 desaturases in PUFA metabolism^58^. Also, the acetylation of histone tightly regulates sperm development, especially during the late-stage differentiation^59, 60, 61, 62^. During spermiogenesis, matured spermatozoa acquire a highly condensed genome by replacing 95% of histone into protamine, a small protein facilitate chromatin condensation. The disruption of histone acetylation impairs histone replacement and causes defects in sperm development^59, 63^. It will be interesting to investigate how IDH3B regulates these biosynthesis pathways and acetylation status in future studies.

In conclusion, our studies show a tissue-specific requirement of IDH3B on modulating mitochondrial metabolism and tissue functions. IDH3B is required for testis metabolism and proper sperm development.

## Materials and Method

### All the chemicals and reagents are detailed in the Key Resource Table (Table S2)

#### Animals

*Idh3b* heterozygous (HET) male mice were purchased from the Jackson Laboratory *(stock #: 042302-JAX)* and crossed with wild-type female mice (C57BL/6J) to transmit the Idh3b-deficient allele to their offspring. The HET males were bred with *Idh3b* homozygous knockout (KO) females to generate *Idh3b*-KO mice and HET littermate controls for experiments. Mouse genotyping was accomplished by Transnetyx, Inc (Cordova, TN, USA) using provided primers (see details in **Table S1**). Both sexes were used in visual studies, but only males for sperm studies. The mouse experiments were performed in accordance with the National Institutes of Health guidelines and the protocol approved by the Institutional Animal Care and Use Committee of West Virginia University.

#### Western blot

The tissues were lysed in RIPA buffer supplemented with protease and phosphatase inhibitor at 5mg/ml concentration. The supernatant containing proteins was collected after centrifuging tissue lysates at 12,000 rpm in 4°C. Protein concentration was determined by the BCA protein assay kit, and 20μg protein samples were boiled and loaded onto SDS-PAGE gels. The gels were transferred to 0.22um nitrocellulose membranes and blocked with 5% non-fat milk in 1X Tris-Buffered Saline containing 0.1% Tween® 20 (TBST). The membranes were incubated with primary antibodies (See details in **Table S2**) at 4°C overnight. After three washes (15min, 5min, 5min) with 1×PBS containing 0.1% Tween-20 (PBST), the membranes were incubated with a rabbit or mouse secondary antibody conjugated with horseradish peroxidase (1:2000) or Alexa Fluor 680 (1: 10,000) for 1 hour, followed by three washes (15min, 5min, 5min) with PBST. A chemiluminescence reagent kit was used to visualize protein bands with horseradish peroxidase secondary antibodies. Western blot using Alexa Fluor secondary antibody were performed as previously reported^64^.

#### Enzyme activity measurement

Samples were harvested from mice at P50 and homogenized in 50mM HEPES/KOH buffer (pH 7.4) containing 1%triton, 1mM EDTA, 10% glycerol and 5mg/ml protease and phosphatase inhibitor. After centrifuged at 12,000 rpm at 4°C, the supernatants were assayed for protein concentrations, and 50ug proteins were loaded for enzymatic activity assays. IDH3 activity was determined by incubating the protein extracts in the assay buffer containing 2mM Mn^2+^, 2mM NAD, 10mM isocitrate at 37 □ in a plate reader and monitoring the production of NADH continuously for 4 hours at 340nm wavelength. IDH1&2 activities were determined by monitoring the production of NADPH at 340nm wavelength for 4 hours after incubation with assay buffer containing 1mM Mg^2+^, 1mM NADP and different concentrations of isocitrate (0mM, 1mM, 2mM, 3mM, 4mM and 10mM) at 37□. No enzyme controls were used as blank controls for each sample. Standard curves of NADH and NADPH were measured simultaneously for calculating the concentration of NADH and NADPH.

#### OCT

OCT was conducted as previously described^64^. Mice were anesthetized with 1.5% isoflurane in a chamber and then received constant isoflurane during OCT via a nose cone. Both pupils were diluted before OCT measurement by applying one drop of 1:1 mixture of 2.5% phenylephrine hydrochloride and 1% tropicamide. To avoid cataract, each eye was applied a drop of GenTeal® lubricant eye gel (Alcon, Fort Worth, TX) right after the dilation, and the body temperature of mice was maintained by a heating platform. Cross-sectional images of the retina were taken using the Bioptigen R-series spectral domain ophthalmic imaging system (Bioptigen, Inc.), and the thickness of each layer was analyzed as previously described ^64, 65^.

#### ERG

Mice were dark-adapted overnight before recording ERG using Celeris D430 rodent ERG testing system (Diagnosys). Under dim red-light illumination, mice were anesthetized, and eyes were dilated as previously described^64^. Scotopic recordings were elicited using light flashes at increasing intensities (0.004, 0.01, 0.063, 0.16, 0.36, 0.8, 1.26, 4, 10 cd.s/m^2^). The photopic response was elicited with increasing flash intensities (1.26, 4, 10, 20 cd.s/m^2^) after a short light adaptation with continuous background light.

#### H&E staining and analysis

The preparation and staining for eye, testis and cauda epididymis were performed as reported before^64^. The tissues were fixed, paraffin sectioned and stained with hematoxylin and eosin. The slides were imaged under a Nikon C2 confocal microscope, and the thickness of outer segment and inner segment in the retina was measured by Image J at 6 different positions of the retina as described before^64^. For testis, 20 seminiferous tubules at four quadrants of testes sections were randomly selected and quantified for the number of sperm cells at different developmental stages.

#### PAS staining and analysis

Similar to H&E staining, testes were embedded in paraffin and sectioned at 4-6 microns. The sections were stained with 1% Periodic Acid, McManus Schiff Stain Reagent, and Gill III Hematoxylin. The stained sections were imaged by a confocal microscope and staged for sperm differentiation based on acrosomal morphology.

#### Sperm cell video recording and counting

For sperm cell video, we dissected cauda epididymis in 1 ml α Minimum essential medium and recorded the sperm movement using EVOS FL Auto 2 Cell Imaging System. To count the number of sperm cells, we sacrificed the mice and dissected the cauda epididymis in 1ml PBS and spun the mix at 1500 RPM for 3 minutes. The aggregates were fixed in 4% paraformaldehyde, resuspended in PBS and counted using a hemocytometer.

#### Immunofluorescent

Testes were immediately fixed in 4% paraformaldehyde for 24 hours at 4□ and rinsed by PBS, dehydrated by 20% sucrose, and then cryopreserved in optimal cutting temperature compound to section at 10um for immunostaining. Sections were washed with PBS containing 1% Triton X-100 for three times and incubated in blocking buffer (10% goat serum with 0.5% triton) at room temperature (RT) for an hour before staining with antibodies or dyes. Fluorescent sections were imaged by a Nikon C2 confocal microscope system.

#### TEM

Testes were cut into half and fixed in 4% glutaraldehyde (Electron Microscopy Sciences) for 30 minutes, then sliced into ~1mm^3^ pieces and continued to be fixed in the same buffer overnight at 4°C. The prefixed testes were washed 5 x 5 minutes in 0.1M cacodylate buffer and postfixed in osmium ferrocyanide for 1 hour on ice. Next, the testes were incubated in a 1% thiocarbohydrazide solution for 20 minutes following by 2% osmium tetroxide for 30 minutes at RT. The tissues were en block stained in 1% uranyl acetate, (aqueous), overnight in the refrigerator and en bloc stained in Walton’s lead aspartate for 30 minutes at 60 °C the second day. Then, the testes were dehydrated in ice-cold 30%, 50%, 70%, and 95% ETOH and infiltrated in a 1:1 mixture of propylene oxide: Durcupan resin, for 2 hours and then overnight infiltration in fresh Durcupan. The next day the testes were placed in fresh Durcupan for two hours, positioned in flat embedding molds and polymerized in a 60 °C oven for two days. 80nm sections were cut on a Leica EM UC7 ultramicrotome and sections were viewed on a JEOL 1230 TEM at 80KV.

#### Steady state metabolomics and ^13^C labeled metabolite analysis by GC MS

Steady state metabolomics and ^13^C labeled metabolite analysis for retina were performed as described in detail before^64^. The metabolites were extracted in 80% cold methanol (V/V), dried and derivatized by methoxymine hydrochloride followed by Ntertbutyldimethylsilyl-N-methyltrifluoroacetamide (TBDMS). An Agilent 7890B/5977B GC/MS was used for GC separation and analysis of metabolites. The data was analyzed by Agilent MassHunter Quantitative Analysis Software and natural abundance was corrected by ISOCOR software.

#### Targeted metabolomics by LC MS/MS

Targeted metabolomics by LC MS/ was performed as reported before^64^. Briefly, metabolites from testes were extracted in 80% cold methanol and analyzed by a Shimadzu LC Nexera X2 UHPLC coupled with a QTRAP 5500 LC MS/MS (AB Sciex) in multiple reaction monitoring (MRM) mode. Each metabolite was tuned with standards for optimal transitions (See details in table S3). The extracted MRM peaks were integrated using MultiQuant 3.0.2 software (AB Sciex).

#### Statistics

The significance of differences between means was determined by unpaired two-tailed T tests using Prism 9.0. Only P < 0.05 was considered to be significant. All data are presented as the mean□±□SD. Multivariate analysis of metabolomics data was performed with a supervised classification model partial least-squares discriminant analysis (PLS-DA) after pareto scaling using MetaboAnalyst 5.0 (https://www.metaboanalyst.ca/). Volcano plot (P<0.05 and fold change>1.5) was performed to analyze the changes of specific metabolites using MetaboAnalyst 5.0.

## Supporting information

Video1 Live sperms in Idh3b HET mice

Video2 Live sperms from Idh3b KO mice

## Acknowledgments

This work was supported by National Institutes of Health Grant EY031324 (JD), EY032462(JD), Bright Focus Foundation M2020141(JD), the Retina Research Foundation (JD) and funds for Core facilities P20 GM103434 (WV INBRE grant), WVCTSI grant GM104942.

We thank Dr. Kolandaivelu for the GM130 antibody and Dr. Ramamurthy for Acetyl-tubulin antibody.

## Ethical approval

The mouse experiments were performed in accordance with the National Institutes of Health guidelines and the protocol approved by the Institutional Animal Care and Use Committee of West Virginia University.

## Informed consent

The authors declare no patients involved in this study and no informed consent was made.

## Data Availability Statement

GC MS and LC MS data generated in this study were provided in the supplemental material. Source data are provided with this paper.

## Author Contributions

Conceptualization J.D and S.Z; Investigation, J.H., R.X., Y.W., Y.W., R.M., E.P., D.K., M.Y., C.Z., B.W., and J.D.; Writing, S.Z., and J.D.; Funding Acquisition, J.D; Supervision, J.D.

## Abbreviations

IDH3B: isocitrate dehydrogenase3 β subunit
IDH3A: isocitrate dehydrogenase3 α subunit
IDH3G: isocitrate dehydrogenase3 γ subunit
IDH1: isocitrate dehydrogenase1
IDH2: isocitrate dehydrogenase2
TCA cycle: Tricarboxylic acid cycle
αKG: α-ketoglutarate
GC MS: Gas Chromatography Mass Spectrometry
LC MS: Liquid Chromatography Mass Spectrometry
OCT: Optical Coherent Tomography
ERG: electroretinogram
IF: Immunofluore scent
WB: Western Blot
HET: heterozygous
KO: knockout
αKG: α-ketoglutarate

