## Supplementary information for "Isocitrate dehydrogenase 3b is required for spermiogenesis but dispensable for retinal degeneration"

This PDF file includes:

Supplementary Figures 1-5

Supplementary Table 1-3

**Supplementary Figures**

**Supplementary Figure 1**

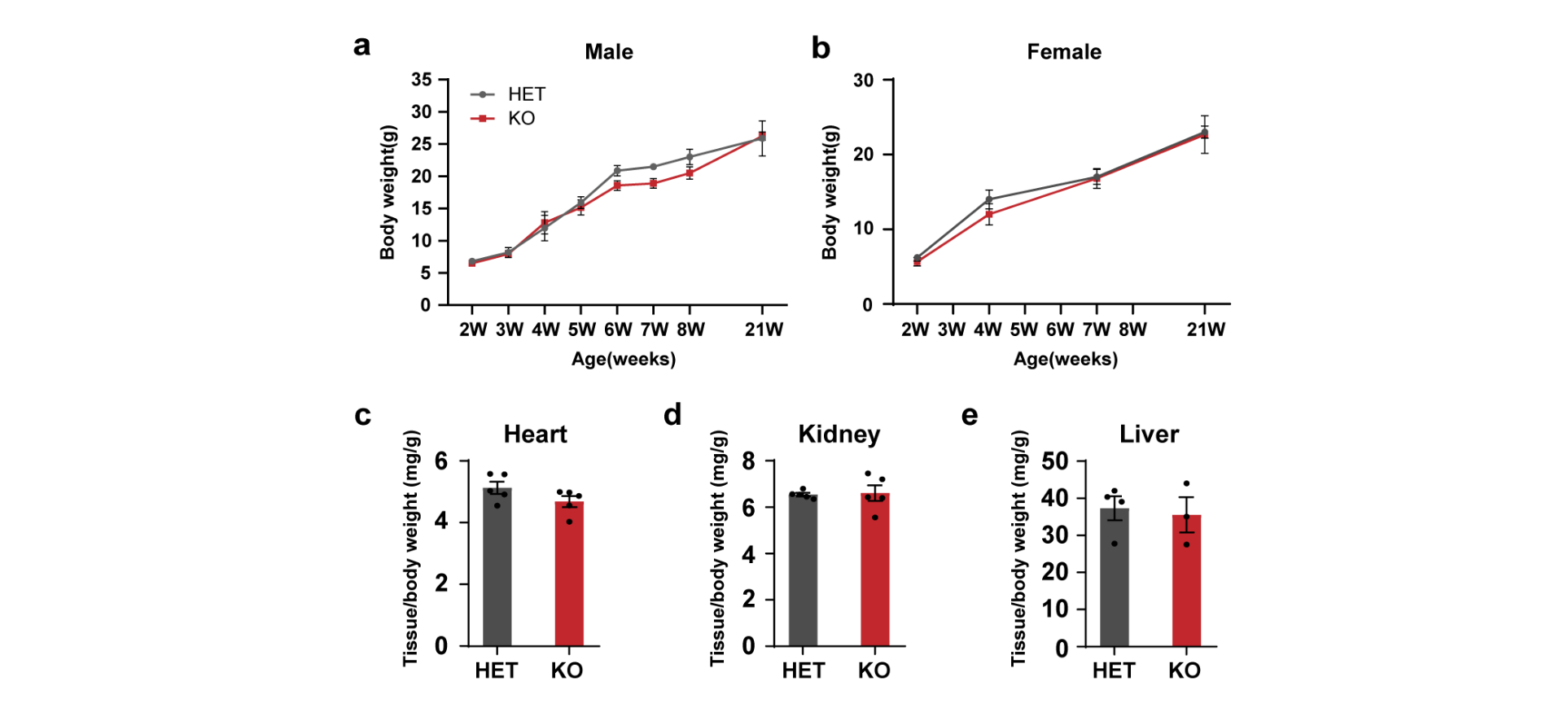

**Supplementary Figure 1. *Idh3b*-KO mice have normal body growth. a-b** Body weight of male and female mice from 2 weeks old to 21 weeks old. N ≥ 4. **c-e** Tissue over body weight ratio for heart, kidney, and liver from mice from P50 to P80. N ≥ 3. (Data are all represented as mean ± SD. t test.)

**Supplementary Figure 2**

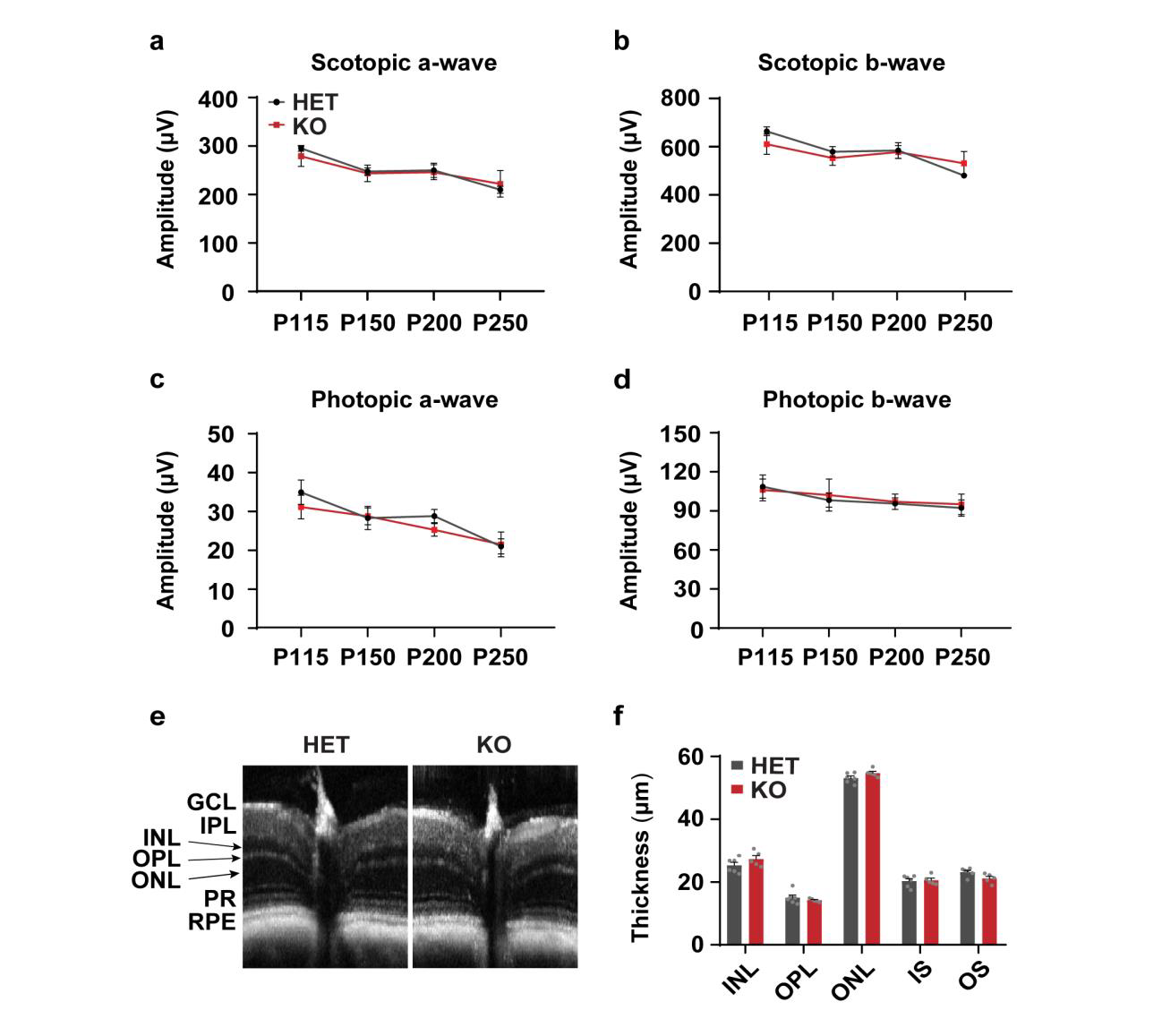

**Supplementary Figure 2. The visual function and retinal morphology are normal in *Idh3b-*KO mice. a-d** Representative scotopic (0.8 cd*s/m2) and photopic (10 cd*s/m2) ERG traces at P115, P150, P200 and P250. N ≥ 4. **e-f** Representative retinal OCT images from HET and KO mice at P200 (e) and the quantitation of thickness of different retinal layers (f). Inner nuclei layer (INL); outer plexiform layer (OPL); outer nuclei layer (ONL); inner segment (IS); and outer segment (OS). N ≥ 5. (Data are all represented as mean ± SD. t test.)

**Supplementary Figure 3**

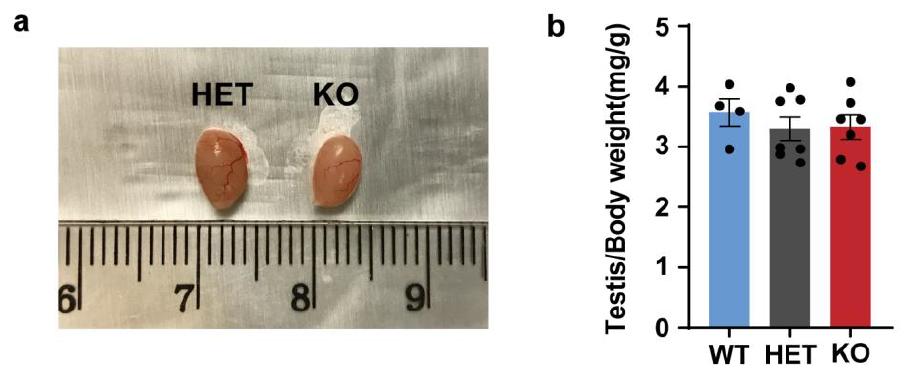
**Supplementary Figure 3. *Idh3b*-KO mice have normal size and weight of their testes. a** Images of testes from HET and KO mice at P50. **b** Ratio of testis weight to body weight from mice at P50 to P80. N ≥ 4. Data are represented as mean ± SD. t test.

**Supplementary Figure 4**

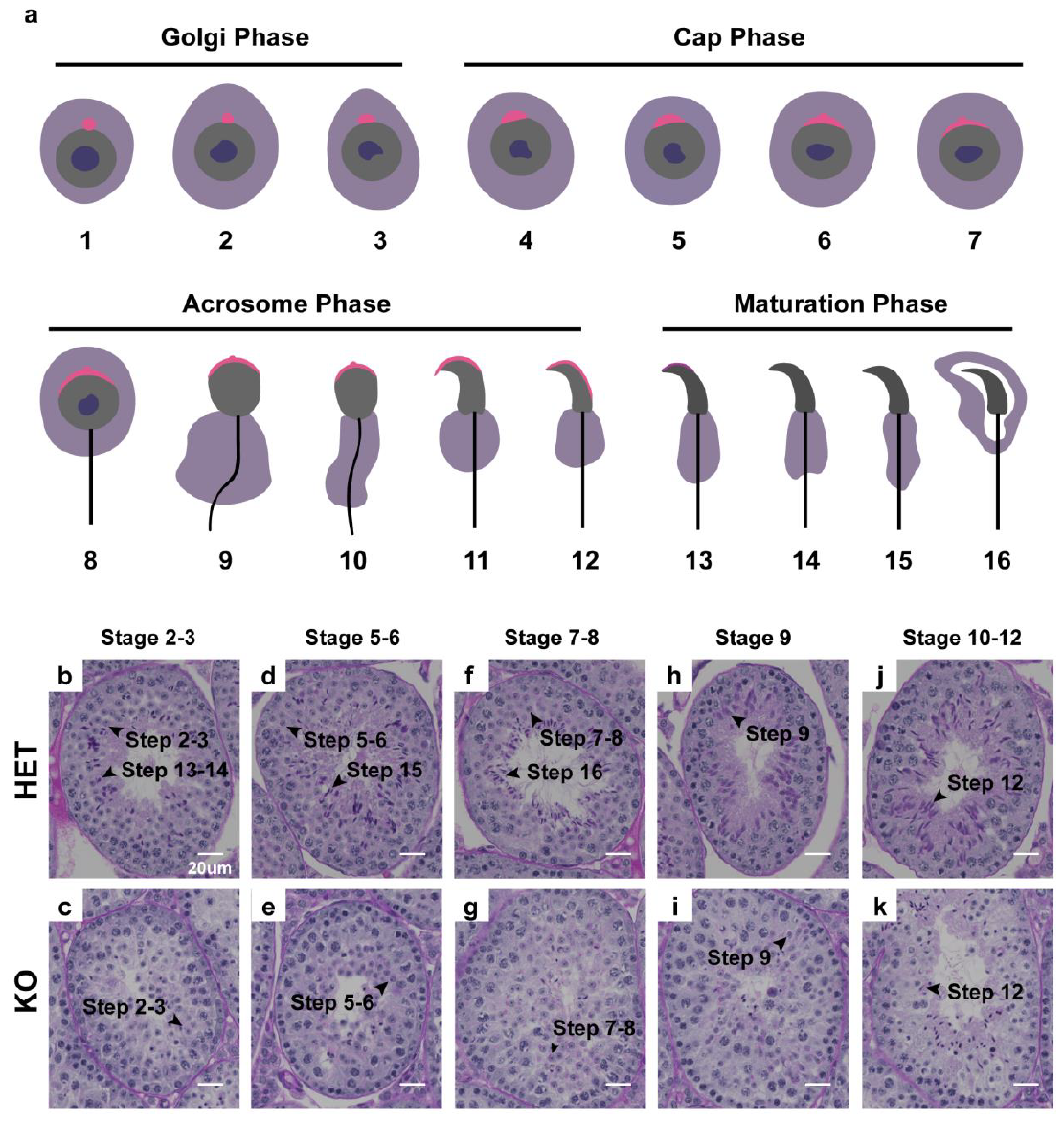

**Supplementary Figure 4. Spermiogenesis is arrested in *Idh3b* KO mice. a** Schematic of 16 steps during spermiogenesis and their corresponding phases for acrosome development. Pink: acrosome; grey: nucleus; dark blue: nucleolus, purple: cytosol; black: flagella. Golgi Phase (steps 1-3): Acrosome originates from a single and sphere acrosomal granule; Cap Phase (steps 4-7): Acrosome develops into a cap-like structure with an acrosomal granule in the center; Acrosome Phase (steps 8-12): Acrosome expanded and nucleus undergo condensation. Maturation Phase (step 13-16): Acrosome forms anterior apex and covers the surface of condensed nucleus with the development of the tail. **b-k** Representative PAS-stained testicular tubules containing spermatids at different differential steps from HET and KO mice at P50. Arrowheads point to spermatids.

**Supplementary Figure 5**

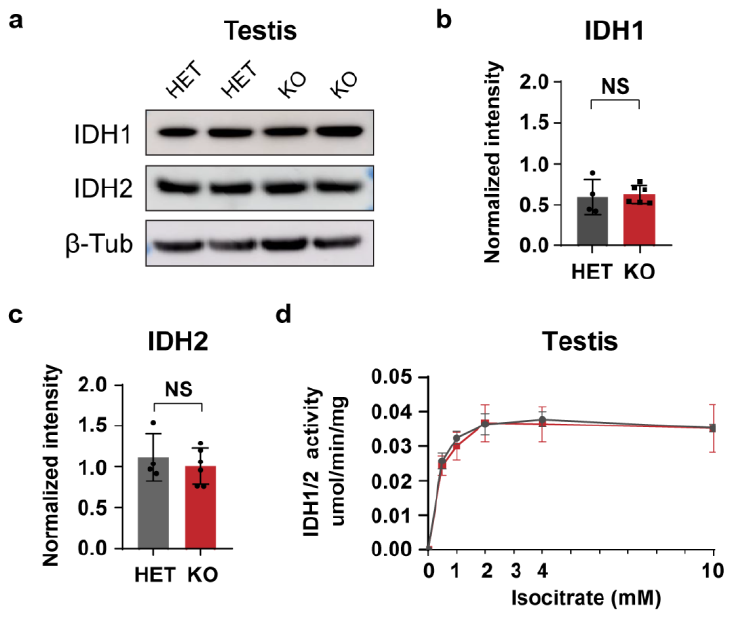

**Supplementary Figure 5. IDH1&2 protein and activities are normal in the testes from Idh3b KO mice. a** Representative immunoblots of IDH1&2 protein expression in testes from mice at P52. **b-c** Relative densitometry of IDH1 and IDH2 levels in testes. β-Tubulin was used as the loading control. N ≥ 4. **d** Enzymatic activity of IDH1&2 under different isocitrate concentrations in testes from HET and KO mice. N ≥ 4. (Data are all represented as mean ± SD. t test.)

**Supplementary Table1. Primers used in experiments**

| Primer IDs | Sequence (5’ to 3’) |
| --- | --- |
| IDH3B-WT For | CTACAGGGATGGATGGGTTGTT |
| IDH3B-WT Rev | CCTTCAGGGATAAGTCCAAGCA |
| IDH3B-KO For | AAGGTCATCGCGGTGTAAGG |
| IDH3B-KO Rev | CAACAACAACAACAACAACAACAAACAA |

**Supplementary Table2. Key resources**

| **Reagents** | **Catalog** | **Company** | **Location** |
| --- | --- | --- | --- |
| RIPA Lysis and Extraction Buffer | 89900 | Thermo scientific | Rockford IL USA |
| Pierce Protease and Phosphatase Inhibitor Mini Tablets | A32959 | Thermo scientific | Rockford IL USA |
| Pierce™ BCA Protein Assay Kit | 23225 | Thermo scientific | Rockford IL USA |
| TGX Stain-Free™ FastCast™ Acrylamide Kit, 10% | 1610183 | Bio-Rad | Hercules CA USA |
| 10x Tris/Glycine/SDS | 1610732 | Bio-Rad | Hercules CA USA |
| 10x Tris/Glycine Buffer for Western Blots and Native Gels | 1610734 | Bio-Rad | Hercules CA USA |
| Nitrocellulose membrane | 1620097 | Bio-Rad | Hercules CA USA |
| Blotting-Grade Blocker (non-fat dry milk) | 1706404 | Bio-Rad | Hercules CA USA |
| Immobilon Western Chemiluminescent HRP Substrate | WBKLS0500 | EMD Millipore Corporation | Burlington MA USA |
| FilterMax™ F5 Multi-Mode Microplate Reader | N/A | Molecular Devices | San Jose CA USA |
| Tween® 20 | 822184 | EMD Millipore Corporation | Billerica MA USA |
| HEPES | H4034-1KG | Sigma-Aldrich | St. Louis MO USA |
| KOH | 8.14353.1000 | Sigma-Aldrich | St. Louis MO USA |
| MnSO4 | 7899-500G | Sigma-Aldrich | St. Louis MO USA |
| NAD | N1511-250MG | Sigma-Aldrich | St. Louis MO USA |
| NADH | N8129-100MG | Sigma-Aldrich | St. Louis MO USA |
| Isocitrate | 205010050 | Acros Organics | Belgium |
| MgCl2 | N7304-100G | Sigma-Aldrich | St. Louis MO USA |
| NADP | 10004675 | Cayman Chemical | Ann Arbor MI USA |
| NADPH | 9000743 | Cayman Chemical | Ann Arbor MI USA |
| Isoflurane | 07-893-1389 | Patterson Veterinary | Greeley CO USA |
| Phenylephrine hydrochloride | NDC 42702-102-15 | Paragon Bioteck, Inc. | Portland OR USA |
| Tropicamide | NDC 61314-355-01 | Sandoz, Inc. | Princeton NJ USA |
| GenTeal® lubricant eye gel | 300650518017 | Alcon laboratories Inc. | Fort Worth TX USA |
| Gill III Hematoxylin | HXGHE3PT | StatLab Medical | Lodi CA USA |
| Alcoholic Eosin | SL98-16 | StatLab Medical | Lodi CA USA |
| Excalibur’s Alcoholic Z-Fix | N/A | Excalibur Pathology | Norman OK USA |
| 1% Periodic Acid | 6316-6X 500ML | EK Industries, Inc. | Joliet IL USA |
| McManus Schiff Stain Reagent | MER 1071 500ML | EK Industries, Inc. | Joliet IL USA |
| α MEM media | 32561-037 | Gibco | Waltham MS USA |
| 16% Paraformaldehyde | 15710 | Electron Microscopy Sciences | Hatfield PA USA |
| Phosphate buffer saline | P3813-10PAK | Sigma-Aldrich | St. Louis MO USA |
| Tissue-Tek optimal cutting temperature compound | 4583 | Sakura Finetek USA Inc | Torrance CA USA |
| Cryostat machine | Leica CM1850 | Leica Biosystems | Lincolnshire IL USA |
| Triton X-100 | LC262801 | LabChem | Zelienople PA USA |
| Methanol | 14262-1L | Honeywell | Morris Plains NJ USA |
| Methoxyamine hydrochloride | 226904-1G | Sigma-Alderich | St. Louis MO USA |
| N-tert-butyldimethylsilyl-N-methyltrifluoroacetamide (TBDMS) | 190500 | Sigma-Alderich | St. Louis MO USA |
| **Antibodies&Host** | **Company** | **Catalog** | **Dilutions** |
| Anti-IDH3A Antibody, rabbit | Sigma-Aldrich | HPA041465 | 1:1000 (WB) |
| Anti-IDH3B Antibody, rabbit | Sigma-Aldrich | HPA049387 | 1:1000 (WB) |
| Anti-IDH3G Antibody, rabbit | Proteintech Group | 25848-1-AP | 1:1000 (WB) |
| Anti-β-Tubulin Antibody, mouse | Santa Cruz | sc-55529 | 1:1000 (WB) |
| Anti-IDH1 Antibody, rabbit | Cell Signaling Technology | 8137S | 1:1000 (WB) |
| Anti-IDH2 Antibody, rabbit | Proteintech Group | 15932-1-AP | 1:1000 (WB) |
| Anti-HSP90B1 Antibody, rat | Santa Cruz | sc-32249 | 1:1000 (WB) |
| Anti-PICK1 Antibody, rabbit | Proteintech Group | 10983-2-AP | 1:1000 (WB) |
| Anti-GOPC Antibody, rabbit | Proteintech Group | 12163-1-AP | 1:1000 (WB) |
| Anti-ACRV1 Antibody, mouse | Santa Cruz | sc-398536 | 1:1000 (WB) |
| Anti-β-Actin Antibody, mouse | Santa Cruz | sc-47778 | 1:1000 (WB) |
| PNA488, N/A | Life Technologies | L21409 | 1:500 (IF) |
| Anti-GM130 Antibody, mouse | BD Biosciences | 610822 | 1:500 (IF) |
| DAPI, N/A | BioRad | 1351303 | 1:500 (IF) |
| Anti-Acetylated α Tubulin Antibody, mouse | Santa Cruz | sc-23950 | 1:1000 (WB&IF) |
| Anti-α-Tubulin Antibody, mouse | Sigma-Aldrich | T6793 | 1:1000 (WB) |
| Anti-ODF2 Antibody, rabbit | Proteintech Group | 12058-1-AP | 1:1000 (WB) |
| Anti-AKAP82 Antibody, mouse | Santa Cruz | sc-135827 | 1:1000 (WB) |
| Mito-tracker, N/A | Life Technologies | M7510 | 1:500 (WB) |
| Goat anti mouse 568, goat | Invitrogen | A10037 | 1:500 (IF) |
| Goat anti mouse 488, goat | Invitrogen | A21202 | 1:500 (IF) |
| Goat anti Rabbit HRP, goat | Cell Signaling Technology | 7074S | 1:2000 (WB) |
| Goat anti Mouse HRP, goat | Abcam | ab131368 | 1:2000 (WB) |
| Goat anti Rat 680, goat | Life Technologies | A-21096 | 1:10,000 (WB) |

**WB: Western Blot; IF, immunofluorescence.**

**Supplementary Table 3. Parameters for metabolites measured in LC MS and GC MS.**

| **Metabolite** | **CAS** | **Mode** | **Q1 Mass (Da)** | **Q3 Mass (Da)** | **DP (Volts)** | **CE (Volts)** | **Platform** |
| --- | --- | --- | --- | --- | --- | --- | --- |
| (R)-3-Hydroxybutyric acid | 300-85-6 | Pos | 105.1 | 45 | 20 | 47 | LC MS |
| 1-Methyladenosine | 15763-06-1 | Pos | 282.1 | 150.1 | 48 | 35 | LC MS |
| 2-Methylbutyroylcarnitine | 31023-25-3 | Pos | 246.2 | 85.1 | 100 | 43 | LC MS |
| 3-Aminoisobutanoic acid | 144-90-1 | Pos | 104.1 | 58 | 55 | 38 | LC MS |
| 3-Hydroxypicolinic acid | 19621-92-2 | Neg | 138.1 | 94.1 | -83 | -19 | LC MS |
| 3-Nitrotyrosine | 621-44-3 | Neg | 225 | 163 | -45 | -19 | LC MS |
| 4-Hydroxyphenylpyruvic acid | 156-39-8 | Neg | 179.1 | 97.1 | -40 | -17 | LC MS |
| 4-Hydroxyproline | 51-35-4 | Pos | 132.1 | 86 | 60 | 17 | LC MS |
| 4-Pyridoxic acid | 82-82-6 | Pos | 182.1 | 138 | -59 | -20 | LC MS |
| 5-Aminopentanoic acid | 660-88-8 | Pos | 118.1 | 55.1 | 57 | 24 | LC MS |
| 6-Phosphogluconic acid | 921-62-0 | Neg | 275 | 79 | -100 | -30 | LC MS |
| Acetyl coenzyme A | 102029-73-2 | Pos | 810.1 | 303.1 | 116 | 46 | LC MS |
| AcetylCholine | 51-84-3 | Pos | 146.1 | 87 | 40 | 20 | LC MS |
| Acetylglycine | 543-24-8 | Pos | 118.1 | 30.2 | 30 | 16 | LC MS |
| Acetylproline | 68-95-1 | Neg | 156 | 114 | -25 | -30 | LC MS |
| Adenine | 73-24-5 | Neg | 134 | 107 | -92 | -25 | LC MS |
| Adenosine | 58-61-7 | Pos | 268.1 | 136 | 95 | 38 | LC MS |
| Adenosine monophosphate | 61-19-8 | Neg | 346 | 134 | -82 | -46 | LC MS |
| Adenosine triphosphate | 987-65-5 | Pos | 507.9 | 136.1 | 14 | 60 | LC MS |
| Adipic acid | 124-04-9 | Neg | 145 | 83 | -50 | -17 | LC MS |
| ADP | 58-64-0 | Neg | 426 | 79.1 | -65 | -84 | LC MS |
| AICAR | 3031-94-5 | Pos | 259.1 | 111 | 85 | 34 | LC MS |
| Aminoadipic acid | 542-32-5 | Neg | 160.1 | 116.1 | -50 | -20 | LC MS |
| Anthranilate | 118-92-3 | Neg | 136 | 92 | -83 | -21 | LC MS |
| Argininosuccinic acid | 2387-71-5 | Pos | 291.1 | 69.9 | 70 | 50 | LC MS |
| Arginosuccinate | 2387-71-5 | Pos | 291.1 | 69.9 | 130 | 65 | LC MS |
| Ascorbic acid | 50-81-7 | Neg | 175 | 87 | -63 | -30 | LC MS |
| Azelaic acid | 123-99-9 | Neg | 187 | 97.1 | -73 | -25 | LC MS |
| Benzoic acid | 65-85-0 | Pos | 123 | 77 | 220 | 31 | LC MS |
| Betaine | 590-46-5 | Pos | 118.1 | 58 | 166 | 56 | LC MS |
| Butyrylcarnitine | 25576-40-3 | Pos | 232.2 | 85.1 | 72 | 30 | LC MS |
| Cadaverine | 462-94-2 | Pos | 103 | 77 | 190 | 25 | LC MS |
| Carbamoyl phosphate | 590-55-6 | Neg | 151 | 108 | -105 | -25 | LC MS |
| Carnosine | 305-84-0 | Pos | 227.1 | 110.1 | 157 | 32 | LC MS |
| CDP | 63-38-7 | Pos | 404 | 112 | 40 | 40 | LC MS |
| Choline | 62-49-7 | Pos | 104.1 | 60.1 | 95 | 37 | LC MS |
| cis-Aconitic acid | 499-12-7 | Neg | 173 | 85 | -37 | -18 | LC MS |
| Citraconic acid | 498-23-7 | Neg | 129 | 85 | -21 | -12 | LC MS |
| Citrulline | 372-75-8 | Neg | 174.1 | 131.1 | -32 | -20 | LC MS |
| CMP | 63-37-6 | Neg | 322 | 79 | -40 | -40 | LC MS |
| Coenzyme A | 55672-92-9 | Pos | 768.1 | 261 | 34 | 47 | LC MS |
| Creatine | 57-00-1 | Pos | 132.1 | 90 | 170 | 17 | LC MS |
| Creatinine | 60-27-5 | Pos | 114 | 44 | 71 | 33 | LC MS |
| CTP | 65-47-4 | Neg | 482 | 159 | -40 | -40 | LC MS |
| Cyclic adenosine diphosphate-ribose | 119340-53-3 | Pos | 542 | 136 | 75 | 51 | LC MS |
| Cyclic AMP | 60-92-4 | Neg | 328 | 134 | -87 | -32 | LC MS |
| Cyclic GMP | 7665-99-8 | Neg | 344 | 150 | -37 | -30 | LC MS |
| Cytidine | 65-46-3 | Pos | 244 | 112.1 | 50 | 30 | LC MS |
| Cytosine | 71-30-7 | Pos | 112 | 95 | 97 | 23 | LC MS |
| D-2-Hydroxyglutaric acid | 103404-90-6 | Pos | 149.1 | 77 | 100 | 43 | LC MS |
| Decanoyl-L-carnitine | 3992-45-8 | Pos | 316.3 | 85.1 | 70 | 25 | LC MS |
| Dehydroascorbic acid | 490-83-5 | Pos | 175 | 63.1 | 130 | 12 | LC MS |
| D-Erythrose 4-phosphate | 585-18-2 | Neg | 199 | 97 | -160 | -15 | LC MS |
| D-Fructose | 57-48-7 | Neg | 179.1 | 71 | -120 | -28 | LC MS |
| D-Glucose | 492-62-6 | Neg | 179 | 89 | -50 | -15 | LC MS |
| Dihydroxyacetone phosphate | 57-04-5 | Neg | 169 | 79 | -38 | 37 | LC MS |
| Dopamine | 51-61-6 | Pos | 154 | 91 | 40 | 36 | LC MS |
| D-Ribulose 5-phosphate | 551-85-9 | Neg | 229 | 79 | -70 | -45 | LC MS |
| dUMP | 964-26-1 | Neg | 307 | 79 | -68 | -66 | LC MS |
| Epinephrine | 51-43-4 | Pos | 184.1 | 77.1 | 50 | 52 | LC MS |
| Erythritol | 149-32-6 | Pos | 123 | 80 | 40 | 33 | LC MS |
| FAD | 146-14-5 | Neg | 784.1 | 437.1 | 52 | 43 | LC MS |
| Flavin Mononucleotide | 6184-17-4 | Neg | 455.1 | 243.1 | 14 | 46 | LC MS |
| Folic acid | 59-30-3 | Neg | 440.1 | 311 | -132 | -30 | LC MS |
| Fructose 1,6-bisphosphate | 488-69-7 | Neg | 339.1 | 79 | -56 | -75 | LC MS |
| Gamma-Aminobutyric acid | 56-12-2 | Pos | 104.1 | 87 | 57 | 13 | LC MS |
| GDP | 146-91-8 | Neg | 442 | 79 | -41 | -61 | LC MS |
| Geranyl-PP | 763-10-0 | Neg | 313.1 | 185.1 | -58 | -45 | LC MS |
| Glucoronate | 12/3/6556 | Neg | 193 | 73 | -56 | -20 | LC MS |
| Glucosamine | 3416-24-8 | Neg | 258 | 79 | -80 | -40 | LC MS |
| Glucose 1-phosphate | 59-56-3 | Pos | 259 | 79 | -85 | -60 | LC MS |
| Glucose 6-phosphate | 56-73-5 | Neg | 259 | 79 | -46 | -62 | LC MS |
| Glutamax | 39537-23-0 | Pos | 218.1 | 84 | 50 | 20 | LC MS |
| Glutaric acid | 110-94-1 | Neg | 131 | 87 | -45 | -16 | LC MS |
| Glutathione | 70-18-8 | Neg | 306 | 143.1 | -61 | -26 | LC MS |
| Glyceraldehyde 3-phosphate | 142-10-9 | Neg | 171 | 79 | -74 | -28 | LC MS |
| Glyceric acid | 14028-62-7 | Pos | 107 | 45 | 66 | 16 | LC MS |
| Glycerol | 56-81-5 | Pos | 93 | 51 | 160 | 36 | LC MS |
| Glycine | 56-40-6 | Pos | 76 | 30.2 | 30 | 16 | LC MS |
| Glyoxylic acid | 298-12-4 | Pos | 75 | 57 | 100 | 17 | LC MS |
| Guanine | 73-40-5 | Pos | 152 | 110 | 20 | 28 | LC MS |
| Guanosine | 118-00-3 | Neg | 282.1 | 150 | -67 | -33 | LC MS |
| Guanosine monophosphate | 85-32-5 | Neg | 362.1 | 79 | -41 | -61 | LC MS |
| Guanosine triphosphate | 86-01-1 | Neg | 521.9 | 159 | -39 | -32 | LC MS |
| Heptadecanoic acid | 506-12-7 | Neg | 269.1 | 135.1 | -77 | -35 | LC MS |
| Hexanoylcarnitine | 22671-29-0 | Pos | 260.2 | 85.1 | 80 | 25 | LC MS |
| Hippuric acid | 495-69-2 | Neg | 178 | 77.1 | -61 | -23 | LC MS |
| Histamine | 51-45-6 | Pos | 112 | 95 | 65 | 18 | LC MS |
| Homocysteine | 6027-13-0 | Pos | 136 | 91 | 135 | 24 | LC MS |
| Hydroxykynurenine | 606-14-4 | Neg | 223 | 75 | -150 | -55 | LC MS |
| Hypotaurine | 300-84-5 | Neg | 108.1 | 64 | -40 | -17 | LC MS |
| Hypoxanthine | 68-94-0 | Neg | 135 | 65 | -108 | -37 | LC MS |
| Inosine | 58-63-9 | Neg | 267 | 135 | -123 | -30 | LC MS |
| Inosinic acid | 131-99-7 | Neg | 347 | 79 | -118 | -86 | LC MS |
| Isopentenyl pyrophosphate | 358-71-4 | Neg | 245.1 | 79 | -60 | -55 | LC MS |
| Isobutyryl-L-carnitine | 25518-49-4 | Pos | 232.1 | 85.1 | 70 | 30 | LC MS |
| L-Acetylcarnitine | 14992-62-2 | Pos | 204.1 | 85 | 71 | 30 | LC MS |
| Lactate | 50-21-5 | Neg | 89 | 43 | -60 | -17 | LC MS |
| L-Alanine | 56-41-7 | Pos | 90 | 44 | 50 | 20 | LC MS |
| L-Arginine | 74-79-3 | Pos | 175.1 | 70.1 | 45 | 33 | LC MS |
| L-Asparagine | 70-47-3 | Pos | 133.1 | 70.1 | 55 | 24 | LC MS |
| L-Aspartic acid | 56-84-8 | Pos | 134.1 | 74 | 50 | 20 | LC MS |
| L-Carnitine | 541-15-1 | Pos | 163.1 | 85 | 100 | 15 | LC MS |
| L-Cystine | 56-89-3 | Neg | 239 | 74 | -50 | -25 | LC MS |
| L-Glutamic acid | 56-86-0 | Pos | 148.1 | 84.1 | 51 | 20 | LC MS |
| L-Glutamine | 56-85-9 | Pos | 147.1 | 84.1 | 45 | 23 | LC MS |
| L-Histidine | 332-80-9 | Pos | 156.1 | 110 | 50 | 22 | LC MS |
| L-Homoserine | 1927-25-9 | Pos | 120.1 | 74 | 37 | 16 | LC MS |
| Linoleic acid | 60-33-3 | Pos | 281.1 | 221 | 60 | 13 | LC MS |
| L-Kynurenine | 343-65-7 | Neg | 207.1 | 144 | -58 | -33 | LC MS |
| L-Leucine | 61-90-5 | Pos | 132.1 | 86 | 60 | 16 | LC MS |
| L-Lysine | 56-87-1 | Pos | 147.1 | 84 | 38 | 22 | LC MS |
| L-Malic acid | 6915-15-7 | Neg | 133 | 115 | -50 | -14 | LC MS |
| L-Methionine | 63-68-3 | Pos | 150.1 | 61 | 27 | 43 | LC MS |
| L-Octanoylcarnitine | 25243-95-2 | Pos | 288.3 | 85.1 | 100 | 25 | LC MS |
| L-Palmitoylcarnitine | 1985-18-8 | Pos | 400.4 | 85.1 | 30 | 40 | LC MS |
| L-Phenylalanine | 63-91-2 | Pos | 166.1 | 120 | 50 | 18 | LC MS |
| L-Proline | 147-85-3 | Pos | 116.1 | 70.1 | 70 | 44 | LC MS |
| L-Serine | 56-45-1 | Pos | 106 | 60 | 40 | 22 | LC MS |
| L-Threonine | 72-19-5 | Pos | 120.1 | 102 | 50 | 10 | LC MS |
| L-Tryptophan | 73-22-3 | Pos | 205 | 146 | 75 | 35 | LC MS |
| L-Tyrosine | 60-18-4 | Pos | 182.1 | 136 | 40 | 17 | LC MS |
| L-Valine | 72-18-4 | Pos | 118.1 | 72 | 60 | 14 | LC MS |
| Maleic acid | 110-16-7 | Neg | 115 | 71 | -34 | -13 | LC MS |
| Malonic acid | 141-82-2 | Neg | 103 | 41 | -50 | -38 | LC MS |
| Malonyl coenzyme A | 108347-84-8 | Pos | 854.1 | 347.1 | 75 | 45 | LC MS |
| Melatonin | 73-31-4 | Neg | 231.1 | 58 | -75 | -37 | LC MS |
| Methylmalonic acid | 516-05-2 | Pos | 119 | 65 | 70 | 37 | LC MS |
| Mevalonic acid | 150-97-0 | Neg | 147.1 | 77 | -50 | -20 | LC MS |
| myo-Inositol | 87-89-8 | Neg | 179 | 87 | -105 | -24 | LC MS |
| Myristoyl-L-carnitine | 25597-07-3 | Pos | 372.4 | 85.1 | 70 | 25 | LC MS |
| N1-Methylnicotinamide | 3106-60-3 | Pos | 138 | 78 | 110 | 37 | LC MS |
| N-Acetylasparagine | 4033-40-3 | Pos | 175.1 | 70.1 | 55 | 24 | LC MS |
| N-Acetylglutamic acid | 1188-37-0 | Pos | 190.1 | 84.1 | 51 | 20 | LC MS |
| N-Acetylglutamic acid | 1188-37-0 | Pos | 190 | 102 | 20 | 23 | LC MS |
| N-Acetyl-L-alanine | 97-69-8 | Pos | 132.1 | 44 | 50 | 20 | LC MS |
| N-Acetyl-L-methionine | 1115-47-5 | Pos | 192.1 | 61 | 27 | 43 | LC MS |
| N-Acetyl-L-phenylalanine | 2018-61-3 | Pos | 208.1 | 120 | 50 | 18 | LC MS |
| N-Acetylputrescine | 5699-41-2 | Pos | 131.1 | 114 | 64 | 30 | LC MS |
| N-Acetylserine | 16354-58-8 | Pos | 148.1 | 60 | 40 | 22 | LC MS |
| N-acetyltryptophan | 87-32-1 | Pos | 247.2 | 146 | 75 | 35 | LC MS |
| NAD | 53-84-9 | Pos | 664 | 136 | 27 | 48 | LC MS |
| NADH | 58-68-4 | Neg | 664 | 397 | -7 | -46 | LC MS |
| NADP | 53-59-8 | Pos | 744 | 136 | 9 | 78 | LC MS |
| NADPH | 2646-71-1 | Neg | 744 | 408 | -35 | -52 | LC MS |
| N-Alpha-acetyllysine | 152473-69-3 | Pos | 189.1 | 84.1 | 50 | 32 | LC MS |
| Niacinamide | 98-92-0 | Pos | 123 | 80 | 136 | 20 | LC MS |
| Nicotinamide D4 | 347841-88-7 | Pos | 127 | 84 | 100 | 44 | LC MS |
| Nicotinamide riboside | 1341-23-7 | Pos | 256.1 | 123 | 68 | 18 | LC MS |
| Nicotinate | 56-67-6 | Neg | 122 | 78 | -105 | -18 | LC MS |
| Nicotinic acid | 59-67-6 | Neg | 122 | 78 | -50 | -15 | LC MS |
| Nicotinic acid mononucleotide | 321-02-8 | Pos | 336.1 | 124 | 75 | 17 | LC MS |
| Nicotinic acid N-oxide | 2398-81-4 | Pos | 139 | 65.1 | 121 | 36 | LC MS |
| Ophthalmic acid | 495-27-2 | Pos | 290.1 | 58 | 139 | 56 | LC MS |
| Oxidized Glutathione | 13081-14-6 | Neg | 611 | 306 | -67 | -34 | LC MS |
| Palmitoleic acid | 373-49-9 | Pos | 255.1 | 195 | 60 | 10 | LC MS |
| Palmityl-CoA | 188174-64-3 | Pos | 1006.3 | 499.5 | 45 | 53 | LC MS |
| p-Aminobenzoic acid | 99-96-7 | Pos | 138.1 | 120 | 63 | 24 | LC MS |
| Pantothenic acid | 137-08-6 | Neg | 218.1 | 71 | -70 | -41 | LC MS |
| Phenylpyruvic acid | 156-06-9 | Neg | 163 | 91 | -38 | -14 | LC MS |
| Phosphocreatine | 19333-65-4 | Neg | 210 | 79 | -42 | -49 | LC MS |
| Phosphoenolpyruvic acid | 138-08-9 | Neg | 167.1 | 79 | -38 | -20 | LC MS |
| Phosphoserine | 407-41-0 | Pos | 186 | 70 | 50 | 26 | LC MS |
| Pipecolic acid | 535-75-1 | Pos | 130 | 84.1 | 58 | 32 | LC MS |
| Propionic acid | 79-09-4 | Pos | 75 | 57 | 29 | 15 | LC MS |
| Propionylcarnitine | 17298-37-2 | Pos | 218.1 | 85.1 | 60 | 24 | LC MS |
| Pyridoxal | 66-72-8 | Neg | 166 | 79 | -30 | -47 | LC MS |
| Pyridoxal 5'-phosphate | 54-47-7 | Neg | 246 | 79.1 | -61 | -61 | LC MS |
| Pyridoxamine | 85-87-0 | Pos | 169.1 | 134.1 | 20 | 36 | LC MS |
| Pyridoxine | 65-23-6 | Pos | 170.1 | 134 | 40 | 29 | LC MS |
| Pyroglutamic acid | 98-79-3 | Pos | 130.1 | 84 | 100 | 20 | LC MS |
| Quinic acid | 77-95-2 | Neg | 191.1 | 85 | -60 | -33 | LC MS |
| Quinolinic acid | 89-00-9 | Neg | 166 | 78 | -53 | -20 | LC MS |
| Retinol | 68-26-8 | Neg | 285.2 | 79 | -79 | -18 | LC MS |
| Riboflavin | 83-88-5 | Pos | 377.1 | 243.1 | 14 | 29 | LC MS |
| S-Adenosyl-L-methionine | 29908-03-0 | Pos | 399.3 | 136 | 55 | 40 | LC MS |
| S-Adenosylmethionine | 86867-01-8 | Pos | 399.3 | 136 | 60 | 30 | LC MS |
| Salicyluric acid | 69-72-7 | Neg | 137 | 93 | -100 | -38 | LC MS |
| Sarcosine | 107-97-1 | Pos | 90 | 44 | 38 | 17 | LC MS |
| Shikimic acid | 138-59-0 | Neg | 173 | 93 | -61 | -23 | LC MS |
| Spermidine | 124-20-9 | Pos | 146.1 | 72 | 60 | 19 | LC MS |
| Spermine | 71-44-3 | Pos | 203.1 | 129 | 33 | 26 | LC MS |
| Sphinganine | 764-22-7 | Pos | 302.3 | 81 | 160 | 47 | LC MS |
| Stearoylcarnitine | 25597-09-5 | Pos | 428.5 | 85.1 | 70 | 25 | LC MS |
| Succinic acid semialdehyde | 692-29-5 | Neg | 101 | 55 | -60 | -45 | LC MS |
| Succinyl-CoA | 108347-97-3 | Pos | 868.1 | 361.2 | 45 | 41 | LC MS |
| Taurine | 107-35-7 | Pos | 126 | 108 | 200 | 15 | LC MS |
| Thiamine | 59-43-8 | pos | 265 | 122.1 | 67 | 40 | LC MS |
| Thiamine Monophosphate | 273724-21-3 | Pos | 345 | 126 | 40 | 36 | LC MS |
| Thiamine Pyrophosphate | 154-87-0 | Pos | 425 | 126 | 40 | 35 | LC MS |
| Trigonelline | 535-83-1 | Pos | 138 | 92 | 60 | 27 | LC MS |
| Tryptamine | 61-54-1 | Pos | 161.1 | 144 | 58 | 15 | LC MS |
| Tyramine | 51-67-2 | Pos | 138 | 77.1 | 58 | 41 | LC MS |
| UDP-Glucosamine | 17479-04-8 | Pos | 608 | 138 | 70 | 62 | LC MS |
| Uridine | 58-96-8 | Pos | 245 | 113 | 23 | 51 | LC MS |
| Uridine diphosphate glucose | 133-89-1 | Pos | 611 | 499 | 40 | 29 | LC MS |
| UTP | 63-39-8 | Neg | 483 | 79 | -40 | -40 | LC MS |
| Vitamin A | 68-26-8 | Pos | 287.1 | 247 | 60 | 16 | LC MS |
| Xanthine | 69-89-6 | Neg | 151 | 108 | -70 | -23 | LC MS |
| Xanthosine | 146-80-5 | Neg | 283.1 | 151 | -92 | -28 | LC MS |
| Xanthurenic acid | 59-00-7 | Neg | 204 | 160 | -67 | -19 | LC MS |
| XMP | 25899-70-1 | Neg | 363 | 79 | -50 | -45 | LC MS |
| α-ketoglutarate | 328-50-7 | Neg | 145 | 101 | -56 | -11 | LC MS |
| β-Nicotinamide adenine dinucleotide | 20111-18-6 | pos | 665 | 136.1 | 95 | 63 | LC MS |
| 2-hydroxyglutarate | ‎2889-31-8 | Pos | 433 | NA | NA | NA | GC MS |
| 3-Phosphoglyceric acid | 80731-10-8 | Pos | 585 | NA | NA | NA | GC MS |
| Acetoacetic acid | 3483-11-2 | Pos | 188 | NA | NA | NA | GC MS |
| Adipic acid | 124-04-9 | Pos | 317 | NA | NA | NA | GC MS |
| Beta alanine | 107-95-9 | Pos | 260.2 | NA | NA | NA | GC MS |
| beta-hydroxybutyrate | 300-85-6 | Pos | 275 | NA | NA | NA | GC MS |
| Biotin | 58-85-5 | Pos | 529.4 | NA | NA | NA | GC MS |
| Cholesterol | 57-88-5 | Pos | 443 | NA | NA | NA | GC MS |
| Citrate | 77-92-9 | Pos | 591 | NA | NA | NA | GC MS |
| Cystathionine | 56-88-2 | Pos | 302 | NA | NA | NA | GC MS |
| Cysteine | 56-89-3 | Pos | 406 | NA | NA | NA | GC MS |
| Fumarate | 110-17-8 | Pos | 287 | NA | NA | NA | GC MS |
| Glycerate | 6000-40-4 | Pos | 391 | NA | NA | NA | GC MS |
| Glycerol | 56-81-5 | Pos | 377 | NA | NA | NA | GC MS |
| Isocitrate | 1637-73-6 | Pos | 591 | NA | NA | NA | GC MS |
| Isoleucine | 73-32-5 | Pos | 200 | NA | NA | NA | GC MS |
| Malate | 6915-15-7 | Pos | 419 | NA | NA | NA | GC MS |
| N-Acetylaspartic acid | 997-55-7 | Pos | 460 | NA | NA | NA | GC MS |
| Ornithine | 70-26-8 | Pos | 286 | NA | NA | NA | GC MS |
| Oxalic acid | 144-62-7 | Pos | 261 | NA | NA | NA | GC MS |
| Oxaloacetate | 328-42-7 | Pos | 332 | NA | NA | NA | GC MS |
| Phosphoenolpyruvic acid | 5541-93-5 | Pos | 453 | NA | NA | NA | GC MS |
| Proline | 147-85-3 | Pos | 184 | NA | NA | NA | GC MS |
| Pyruvate | 113-24-6 | Pos | 174 | NA | NA | NA | GC MS |
| Quinolinic acid | 89-00-9 | Pos | 338 | NA | NA | NA | GC MS |
| Succinate | 110-15-6 | Pos | 289 | NA | NA | NA | GC MS |
| Uracil | 66-22-8 | Pos | 283.3 | NA | NA | NA | GC MS |
| Urea | 57-13-6 | Pos | 231 | NA | NA | NA | GC MS |

**Pos, positive mode; Neg, negative mode; NA, not applicable.**
